# Rationally derived inhibitors of hepatitis C virus (HCV) p7 channel activity reveal prospect for bimodal antiviral therapy

**DOI:** 10.1101/374793

**Authors:** Joseph Shaw, Rajendra Gosein, Monoj Mon Kalita, Toshana L. Foster, Jayakanth Kankanala, D. Ram Mahato, Claire Scott, Barnabas J. King, Emma Brown, Matthew J. Bentham, Laura Wetherill, Abigail Bloy, Adel Samson, Mark Harris, Jamel Mankouri, David Rowlands, Andrew Macdonald, Alexander W. Tarr, Wolfgang B. Fischer, Richard Foster, Stephen Griffin

**Author notes:** Corresponding authors: Dr Stephen Griffin, Leeds Institute of Medical Research, School of Medicine, Faculty of Medicine and Health, and Astbury Centre for Structural Molecular Biology, University of Leeds, Wellcome Trust Brenner Building, St James’ University Hospital, Leeds LS9 7TF., Dr Richard Foster, School of Chemistry, Faculty of Maths and Physical Sciences and Astbury Centre for Structural Molecular Biology, University of Leeds, Leeds, LS2 9JT. Center for Drug Design, University of Minnesota Twin Cities, Nils Hasselmo Hall, 312 Church Street SE, Mail Code 1191, Minneapolis, MN 55455. School of Veterinary Medicine and Science, Faculty of Medicine & Health Sciences, University of Nottingham, Sutton Bonington Campus, Sutton Bonington, Leicestershire, LE12 5RD, United Kingdom (Toshana L. Foster). ReViral Ltd, NETPark Incubator, Thomas Wright Way, NETPark, Sedgefield, County Durham, TS21 3FD, United Kingdom (Claire Scott).

## Abstract

Since the 1960s, a single class of agent has been licensed targeting virus-encoded ion channels, or “viroporins”, contrasting the success of channel blocking drugs in other areas of medicine. Although resistance arose to these prototypic adamantane inhibitors of the influenza A virus (IAV) M2 proton channel, a growing number of clinically and economically important viruses are now recognised to encode essential viroporins providing potential targets for modern drug discovery.

We describe the first rationally designed viroporin inhibitor with a comprehensive structure-activity relationship (SAR). This step-change in understanding not only revealed a second biological function for the p7 viroporin from hepatitis C virus (HCV) during virus entry, but also enabled the synthesis of a labelled tool compound that retained biological activity. Hence, p7 inhibitors (p7i) represent a unique class of HCV antiviral targeting both the spread and establishment of infection, as well as a precedent for future viroporin-targeted drug discovery.

## Introduction

Hepatitis C virus (HCV) represents a global clinical challenge as a major cause of chronic liver disease, with severe complications including cirrhosis, liver failure and primary liver cancers (hepatocellular- and intrahepatic cholangio-carcinomas (HCC, iCCA)). Acute infection is predominantly asymptomatic which, combined with limited awareness and population screening, means that liver disease is often advanced upon diagnosis. WHO estimates put the total number of deaths due to HCV infection in 2015 at more than 400 000, with ∼1.75 million new infections annually.

HCV antiviral therapy, originally comprising recombinant type 1 interferon (IFN) combined with the guanosine analogue ribavirin, has been revolutionised by new direct-acting antivirals (DAA). DAA are an unprecedented drug development success, capable of achieving high rates of cure with favourable toxicity profiles enabling their use in patients with advanced disease(*1*). Current DAA target three proteins within the viral replicase (NS3/4A protease, NS5A and the NS5B RNA-dependent RNA Polymerase (RdRP)), with drug combinations available for treating each of the eight viral genotypes.

However, the absence of an HCV vaccine, or other means of prophylaxis, makes DAA-based eradication strategies proposed for ∼71 million chronically infected individuals immensely challenging. DAA availability remains limited by cost, coincident with poor diagnostic rates and rapidly increasing burden in low/middle income countries (LMIC). Resistant viral variants are an increasing concern(*2*), with recent reports of increased resistance amongst some rarer viral subtypes(*3*). Compliance within high-risk populations is low and successful DAA therapy does not prevent re-infection. Moreover, recent studies suggest that DAA are less able to reduce the risk of HCC in treated patients compared with IFN-based therapy(*4*). This may be linked to virus-induced host epigenetic signatures that are not reversed following DAA cure(*5*).

HCV is an enveloped positive sense RNA virus with a ∼9.6 kb genome encoding a single large polyprotein translated from an internal ribosomal entry site (IRES) in the 5L-untranslated region. The polyprotein is spatially organised into structural components at the amino terminus and replicase proteins towards the carboxyl terminus; these are released by host and viral proteases, respectively. In addition, p7 and NS2 play pivotal roles during virion assembly(*6*) involving protein-protein interactions with one another, as well as other viral proteins(*7–9*). Furthermore, the 63 amino acid p7 protein is capable of oligomerising (forming hexamers and/or heptamers(*10, 11*)) within membranes to form an ion channel complex(*12–15*) with a distinct, but equally essential role during virion secretion(*16, 17*). This comprises the raising of secretory vesicle pH, which is necessary to protect acid-labile intracellular virions(*18–20*).

Prototypic compounds, such as adamantanes and alkyl imino-sugars, inhibit p7 channel activity as well as virion secretion in culture, but with relatively poor potency(*21, 22*). However, identification of explicit resistance polymorphisms confirmed that such effects are specific(*23*). This includes Leu20Phe, which confers resistance in genotype 1b and 2a p7 to adamantanes, including rimantadine. Thus, despite poor potency, prototypic inhibitors highlight druggable regions upon p7 channel complexes suited to targeting by improved compounds.

The majority of p7 structural studies support the folding of protomers into a hairpin conformation(*10, 24–26*). This is in agreement with immuno-gold labelling of p7 channel complexes by electron microscopy (EM)(*10*), immunofluorescence studies of epitope-tagged p7 expressed in mammalian cells(*27*), and the membrane topology necessary to orient NS2 correctly within the ER membrane during translation of the viral polyprotein. Resultant p7 channel models comprise hexa- or heptameric assemblies of tilted protomers and a lumen formed by the N-terminal helix containing a well conserved (∼90 %, see Table S1) His17 residue, as proven biochemically(*28*). However, a solution NMR structure of a complete hexameric p7 channel complex comprised protomers in an unusual intertwined triple helix configuration (PDB: 2M6X)(*29*). This structure retained a wider channel lumen compared to hairpin-based structures and exposed conserved basic residues to the lipid bilayer. Functionality of this sequence was not demonstrable, possibly due to mutagenesis of conserved cysteine residues to enhance recombinant expression. Furthermore, recent studies have questioned the validity of this structure due to potential artefacts caused by alkyl-phosphocholine detergents used as membrane mimetics(*30*); the original authors contest this notion(*31*). However, molecular dynamics simulations favour the stability and channel gating characteristics of hairpin-based structures(*32, 33*).

Interestingly, both hairpin- and triple helix-based channel structures retain an adamantane binding site upon the channel periphery that includes position 20(*24, 29*). Unsurprisingly, the conformation and amino acid content of this site differs significantly between structures. Previously, we used a hairpin-based heptameric channel complex as a template for *in silico* high throughput screening, based upon a genotype 1b monomeric hairpin p7 solution NMR structure (PDB: 3ZD0). Resultant chemical hits displayed considerably improved potency compared with rimantadine that was independent of Leu20Phe mutations(*24*). However, initial hits lacked convergence around a common pharmacophore and this prevented understanding of a structure-activity relationship (SAR).

We now present a second-generation lead-like oxindole based inhibitor of p7 channel activity complete with a comprehensive SAR: “JK3/32”. The resultant step forward in potency and specificity has not only led to the identification of a second biological role for p7 channel activity during virus entry, but also enabled the generation of a modified tool compound that retained biological activity. This distinguishes p7i from other DAAs by targeting two discrete stages of the virus life cycle separate to genome replication and sets a new precedent for viroporin-targeted drug design.

## Results

### Refining a rapid throughput assay for secreted HCV infectivity

p7 channel activity is essential for the secretion of infectious virions(*20*), making secreted infectivity an ideal biomarker readout for inhibitor antiviral effects. To expedite testing of secreted HCV infectivity following treatment with high numbers of compounds and/ or concentration points, we adapted previously published protocols using the IncuCyte ZOOM(*34*) to quantify infection of naïve cells immunostained for NS5A (Fig 1A). Dilution of virus-containing supernatants was optimised (1:4) for signal-to-noise whilst accurately reflecting infectivity (i.e. within the linear range of a dilution series correctly determining virus titre, see Fig S1A). This negates the need for serial dilutions and removes both human error and the amplification of said error due to multiplication by large dilution factors. The assay is applicable to multiple native or chimeric viruses and readily generates 8-point IC_50_ curves in a reporter-free system.

**Figure 1.**
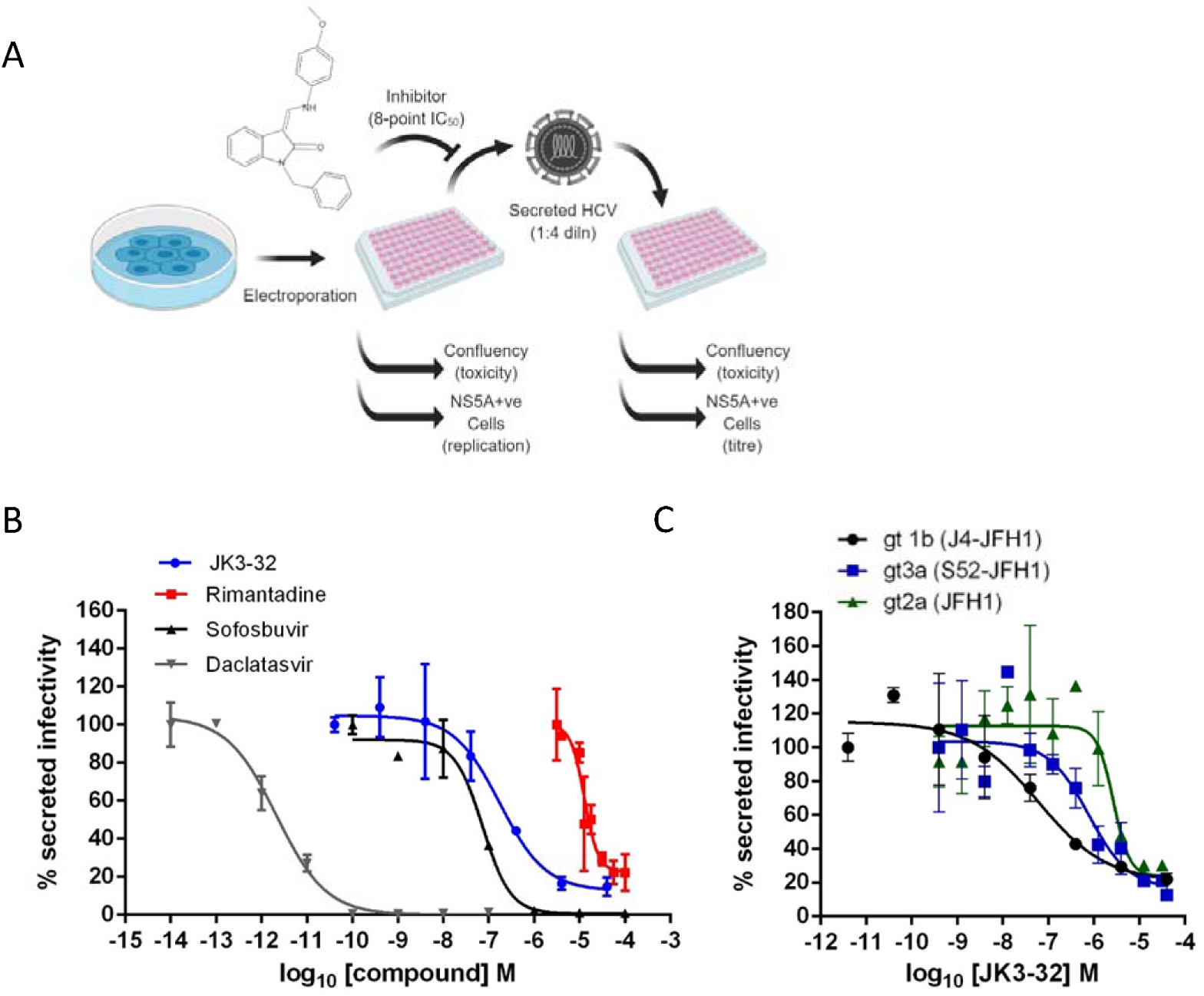
Activity of JK3/32 against HCV particle secretion. **A.** Diagram of workflow for rapid throughput assay for secreted infectivity (generated using “Biorender”, https://app.biorender.com). **B.** Comparison of JK3/32 potency vs. virion secretion of J4/JFH-1 with licensed HCV DAAs Sofosbuvir and Daclatasvir, as well as the prototypic adamantane viroporin inhibitor, rimantadine. Curves are representative of at least four experimental repeats for JK3/32, multiple for Sofosbuvir and Daclatasvir, and two for rimantadine, where each condition is carried out in quadruplicate and error bars represent standard deviations. **C.** Comparative IC_50_ curves for JK3/32 effects upon GT1b, 2a and 3a chimaeric HCV (J4, JFH-1, S52/JFH-1) secreted infectivity post-electroporation. Curves are again representative of multiple experiments and error bars show standard deviations between quadruplicate repeats.

Treatment with the RdRP inhibitor, Sofosbuvir (SOF), reduced viral replication within transfected producer cells (Fig S1B), and this was previously shown to be directly proportional to secreted infectivity(*34*). The NS5A inhibitor, Daclatasvir (DCV) also reduced secreted infectivity, with no effect upon cellular toxicity monitored by producer cell confluency across 8-point dilution ranges (Fig S1C). The ability of the assay to identify false positives caused by cellular toxicity was confirmed using a HSP90 inhibitor, Radicicol, which caused a reduction in secreted infectious particles congruent with producer cell viability (note, cell viability was confirmed by parallel MTT assays, see methods). In addition to detecting anti-viral activity of replication inhibitors SOF and DCV, the assay successfully quantified anti-viral effects of the prototypic p7 inhibitor, Rimantadine, confirming the ability to monitor effects upon p7-dependent virus secretion (Fig 1B). The assay Z factor was determined as 0.47 ± 0.14, with a % coefficient variation of 16.9 ± 4.3 and a signal/ background ratio of 81.1 ± 23.6 (data from 2 independent positive and negative controls (SOF, DCV, DMSO only), in duplicate, averaged over 3 independent experimental repeats, with 3 separate assay plates, over 2 separate days).

### Identification of lead compound JK3/32

SAR-focused chemical modification (Table 1) of an oxindole core scaffold identified “JK3/32” as a series lead with excellent potency against chimaeric genotype 1b HCV (J4/JFH-1) secretion (IC_50_∼184 nM) (Fig 1B, S1C). Thus, JK3/32 potency was comparable to SOF and considerably improved compared with the prototypic p7 inhibitor (p7i), Rimantadine. As seen for first generation compounds(*24*), JK3/32 retained cross-genotype activity versus HCV genotype 3a (IC_50_∼738 nM), with a modest reduction in activity against more genetically distant 2a viruses (IC_50_∼1900 nM) (Fig 1C). The compound showed a toxicity-based selectivity index of >500 (CC_50_ > 100 000 nM based upon confluency (Fig S1C) and MTT assay (Fig S2A) in Huh7 cells, and had no discernible effect upon replication of HCV JFH-1 subgenomic replicons, which retain the same replicase as chimaeric viruses but lack the structural proteins, p7 and NS2 (Fig S2B).

**Table 1.**
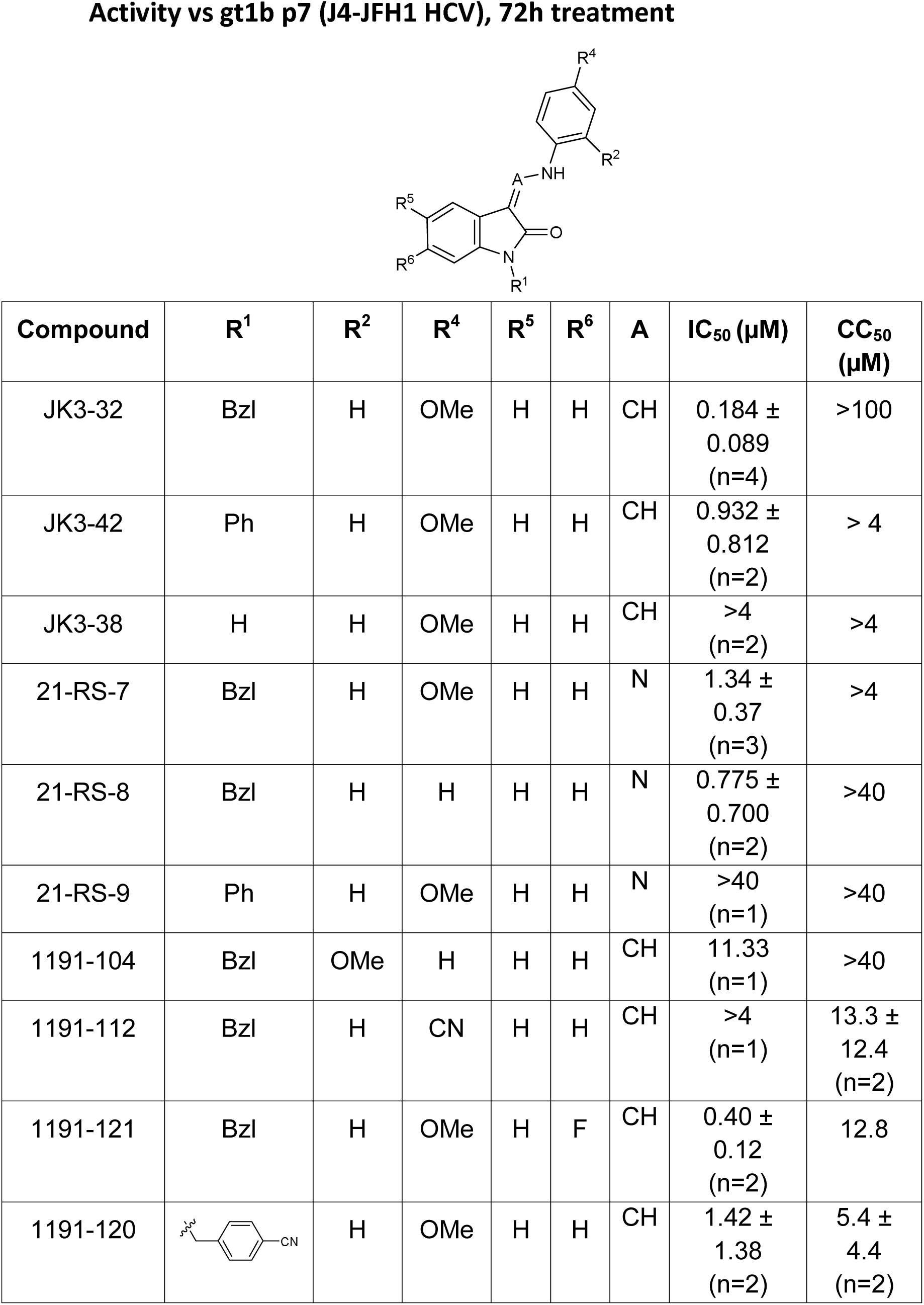
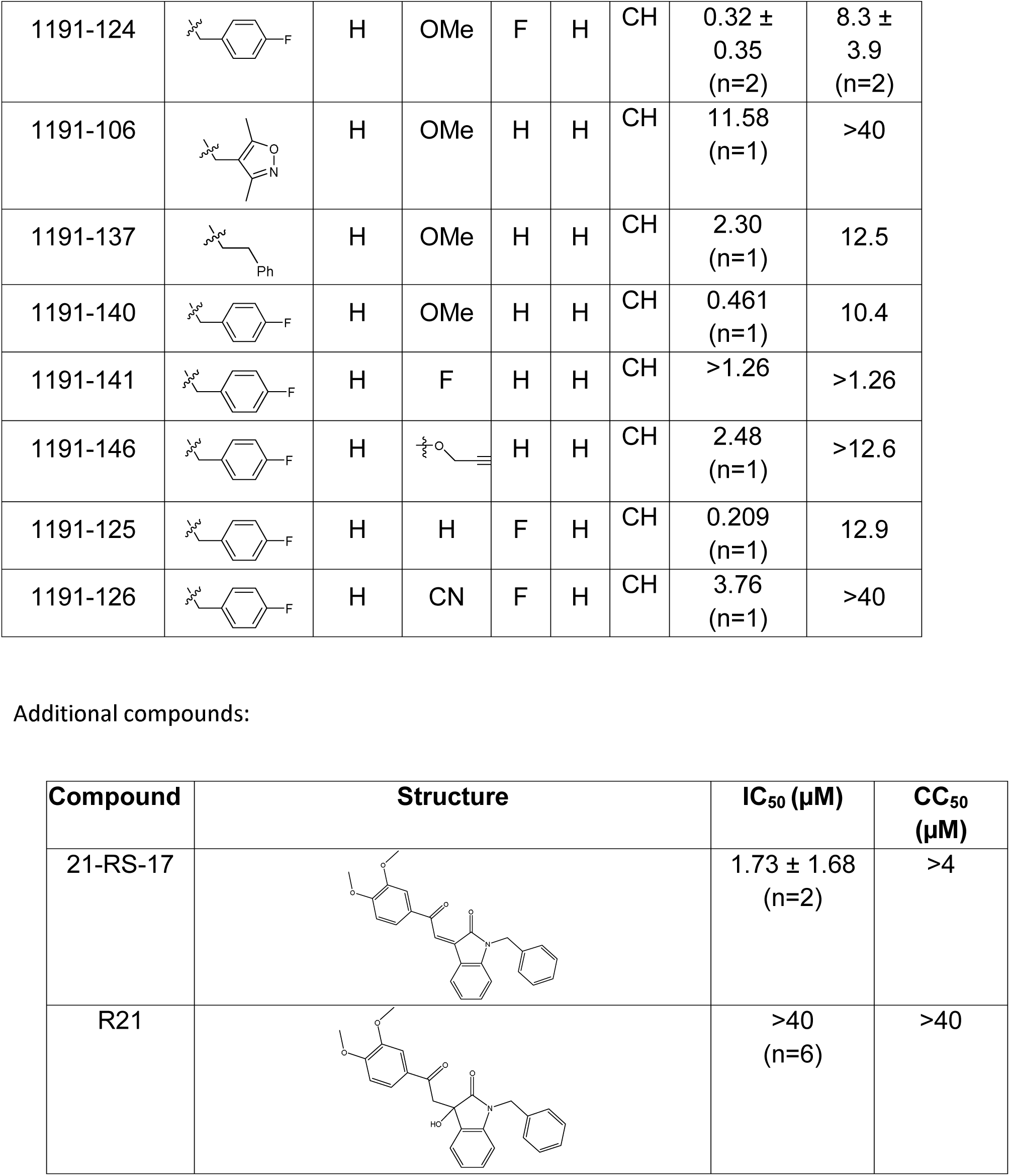
SAR table for JK3/32 series showing compounds contributing directly. Core oxindole scaffold for JK3/32 shown, indicating six R-groups (R^1^-R^6^) subjected to modification, as well as a position within the core structure (A).

| Compound | R<sup>1</sup> | R<sup>2</sup> | R<sup>4</sup> | R<sup>5</sup> | R<sup>6</sup> | A | IC<sub>50</sub> (μM) | CC<sub>50</sub> (μM) |
| --- | --- | --- | --- | --- | --- | --- | --- | --- |
| JK3-32 | Bzl | H | OMe | H | H | CH | 0.184 ± 0.089 (n=4) | >100 |
| JK3-42 | Ph | H | OMe | H | H | CH | 0.932 ± 0.812 (n=2) | > 4 |
| JK3-38 | H | H | OMe | H | H | CH | >4 (n=2) | >4 |
| 21-RS-7 | Bzl | H | OMe | H | H | N | 1.34 ± 0.37 (n=3) | >4 |
| 21-RS-8 | Bzl | H | H | H | H | N | 0.775 ± 0.700 (n=2) | >40 |
| 21-RS-9 | Ph | H | OMe | H | H | N | >40 (n=1) | >40 |
| 1191-104 | Bzl | OMe | H | H | H | CH | 11.33 (n=1) | >40 |
| 1191-112 | Bzl | H | CN | H | H | CH | >4 (n=1) | 13.3 ± 12.4 (n=2) |
| 1191-121 | Bzl | H | OMe | H | F | CH | 0.40 ± 0.12 (n=2) | 12.8 |
| 1191-120 |  | H | OMe | H | H | CH | 1.42 ± 1.38 (n=2) | 5.4 ± 4.4 (n=2) |
| 1191-124 |  | H | OMe | F | H | CH | 0.32 ± 0.35<br>(n=2) | 8.3 ± 3.9<br>(n=2) |
| 1191-106 |  | H | OMe | H | H | CH | 11.58<br>(n=1) | >40 |
| 1191-137 |  | H | OMe | H | H | CH | 2.30<br>(n=1) | 12.5 |
| 1191-140 |  | H | OMe | H | H | CH | 0.461<br>(n=1) | 10.4 |
| 1191-141 |  | H | F | H | H | CH | >1.26 | >1.26 |
| 1191-146 |  | H |  | H | H | CH | 2.48<br>(n=1) | >12.6 |
| 1191-125 |  | H | H | F | H | CH | 0.209<br>(n=1) | 12.9 |
| 1191-126 |  | H | CN | F | H | CH | 3.76<br>(n=1) | >40 |

Additional compounds:
| Compound | Structure | IC<sub>50</sub> (μM) | CC<sub>50</sub> (μM) |
| --- | --- | --- | --- |
| 21-RS-17 |  | 1.73 ± 1.68<br>(n=2) | >4 |
| R21 |  | >40<br>(n=6) | >40 |

The oxindole scaffold of JK3/32 resembles that of certain licensed kinase inhibitor drugs (Sunitinib, Nintedanib). However, the JK series of inhibitors is chemically distinguished from these compounds by an *N*-alkyl substituent, which was essential for anti-HCV activity (Table 1); accordingly, JK3/32 displayed no off-target activity against a panel of human kinases tested commercially (Fig S2C).

JK3/32 was part of a chemical series derived through evolution of an original hit compound, LDS19, selected *in silico* using a 3ZD0-based structure model template(*24*) (Fig S3). The first iteration of compounds (prefix “RS”) was tested using in *vitro dye* release assays(*35*) using genotype 1b p7 (J4 strain) (Fig S4). This confirmed that variation of the prototypic scaffold generated compounds displaying activity versus p7 channel function and that a specific structure-activity relationship (SAR) should be achievable. Cell culture assays confirmed compound activity and comprised the screening method for ensuing compound iterations (Table 1).

Finally, we compared JK3/32 with an amiloride derivative that has been progressed into early phase human trials in Asia. BIT225 was identified as an inhibitor of genotype 1a p7 using a bacterial screen and has been reported to display activity versus Bovine viral diarrhoea virus (BVDV)(*36*), and more recently HCV in cell culture(*37*). However, in our hands BIT225 showed no antiviral activity discernible from effects upon cellular viability (Fig S5); notably, no assessment of cellular toxicity was undertaken during previously reported HCV studies(*37*), which used a concentration higher (30 μM) than the observed Huh7 CC_50_ herein (18.6 μM) during short timescale assays (6-24 h).

### JK3/32 SAR corroborates predicted binding to hairpin-based p7 channel models

We developed a library of JK3/32 analogues to explore SAR for inhibition of J4/JFH-1 secretion (Table 1). Of forty-one compounds tested, twenty contributed directly to the JK3/32 SAR, which was largely consistent with energetically preferred *in silico* docking predictions (using Glide, Schrodinger). JK3/32 is predicted to bind into a predominantly hydrophobic cleft created between helices on the membrane-exposed site (Fig 2a, b). Predicted polar interactions occur between the side-chains of Tyr45 and Trp48 side and the carbonyl oxygen atom at the indole core (Fig 2c). Other predicted close contacts included residues experimentally defined by NMR to interact with rimantadine(*24*): Leu20, Tyr45, Gly46, Trp48, Leu50 and Leu52, and additional interactions with Ala11, Trp32 and Tyr42. Importantly, the majority of residues within this binding site are highly conserved; all residues are >90% conserved with the exception of Leu20 (45.67%) and Tyr45 (84.67%) (Fig 2d, S6, Table S1). However, unlike rimantadine, Leu20Phe does not mediate resistance to this chemical series(*24*).

**Figure 2.**
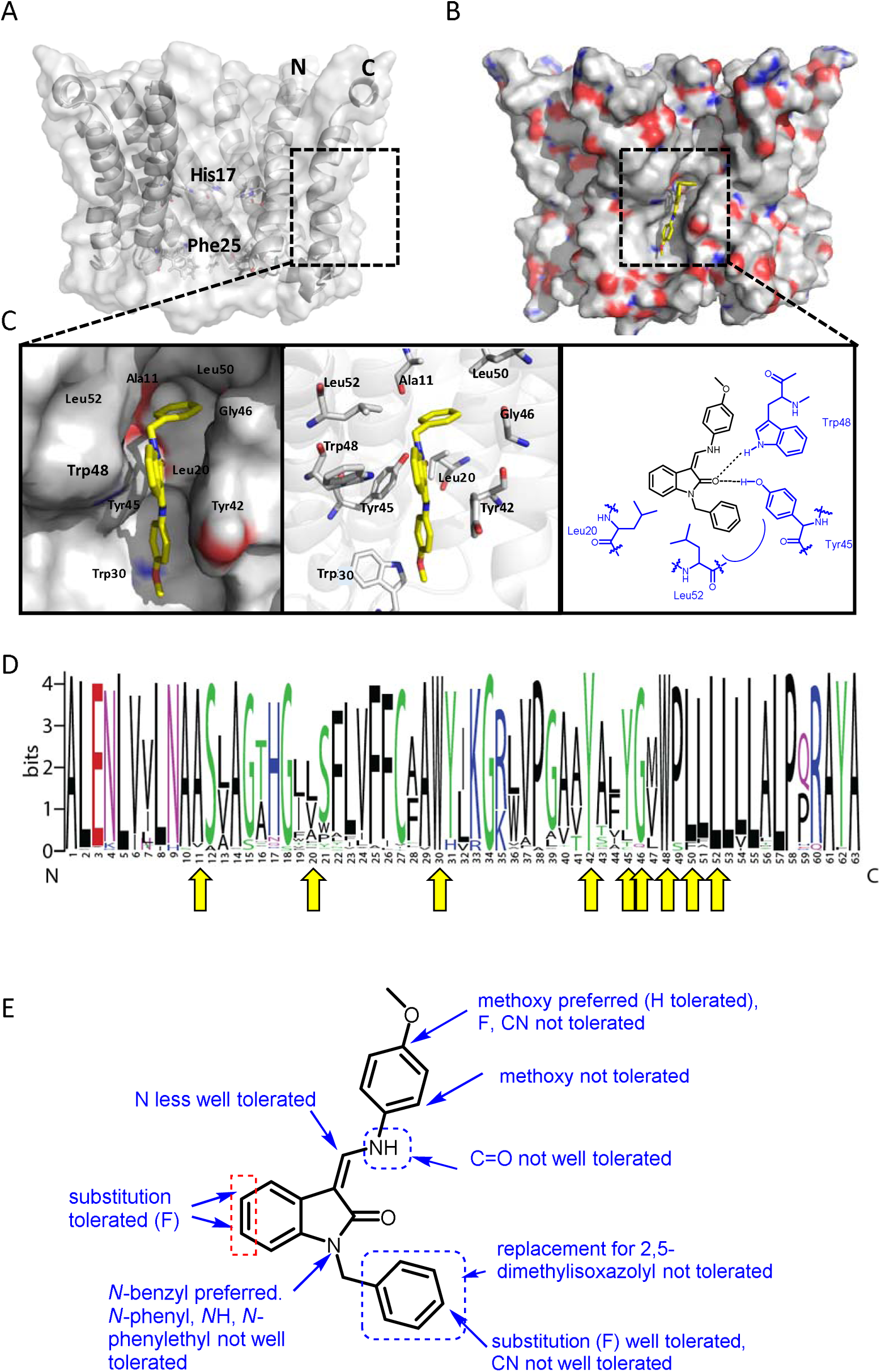
Predicted interactions of JK3/32 with genotype 1b p7 heptamer complexes. **A.** Cutaway image of PDB: 3ZD0-based heptamer showing orientation of N- and C-terminal helices, predicted gating residue (Phe25) and proton sensor (His17). Box shows approximate region corresponding to peripheral drug binding site. **B.** Space-filling model of PDB: 3ZD0-based heptamer showing basic (blue) and acidic (red) charge distribution and positioning of JK3/32 (yellow) within peripheral binding site (box). **C.** Zoomed images showing peripheral drug binding site and predicted energetically preferred binding pose (in Glide) for JK3/32 (yellow) within membrane-exposed binding site as space fill (left), side chains (middle) and key interactions, including with Tyr45 and Trp48 (right). **D.** Amino acid conservation within p7 across ∼1500 sequences from the EU HCV database. Height corresponds to relative conservation of individual residues (quantified in Table S1). Yellow arrows indicate residues predicted to form direct interactions with JK3/32. **E.** Summary of the structure activity relationships for the JK3/32 series of inhibitors (Table 1) for their effects upon chimaeric GT1b HCV (J4/JFH-1) secreted infectivity following electroporation.

JK3/32 SAR was consistent with its predicted binding pose (Fig 2e and Table 1) following docking within the peripheral binding site, defining key determinants of its activity. For example, substitution of the N1 position of the oxindole core demonstrated a preference for benzyl substitution (e.g. JK3/32) consistent with the group occupying a relatively large hydrophobic pocket between helices created by Leu and Ala residues. Incorporation of a longer, more hydrophobic group (N-ethylphenyl, 1191-137), was less well tolerated. Introduction of a NH (JK3-38), N-Ph (JK3-42) and a N-heterocyclic substituent (2,5-dimethylisoxazolylmethyl, 1191-106) abrogated antiviral effects. Attempts at substitution at the ‘northern’ phenyl ring was not well tolerated, with 4*-*OMe (e.g. JK3/32) or H (21-RS-8) preferred over 4-cyano (1191–112) and 2-methoxy (1191–104). 4-alkynyloxy was only moderately less active than 4-OMe (compare 1191-146 (IC_50_ 2.48 μM) to 1191-140 (IC_50_ 0.46 μM)), suggesting that further synthetic expansion from this site was possible, consistent with the modelling which directed this vector outwards from the binding pocket into the membrane. The linker at the 3-position of the oxindole core was sensitive to modification. The hydrazone analogue (21-RS-7) was much less active whilst replacement of the NH for a carbonyl group (21-RS-17) was also not well tolerated. This is consistent with the enamine linker adopting an important bridging unit for correct placement of the N1 substituent into the deep hydrophobic pocket. Substitution of the oxindole core at the 5- and 6-positions with F atoms (1191-124 and 1191-121 respectively) was generally well tolerated. Consistency between observed SAR and the heptameric 3ZD0-based model supports that rational design based upon this system generates authentic, specific p7-targeted antivirals; JK3/32 SAR was not consistent with the 2M6X structure (data not shown).

### Molecular dynamics supports stable JK3/32 binding at the peripheral site

We next undertook atomistic molecular dynamics simulations of genotype 1b p7 channel complexes with JK3/32 bound at the peripheral site to assess the stability of interactions predicted by docking studies over time. Encouragingly, atomistic simulations (100 ns) of JK3/32 bound to the peripheral site revealed marked stability of its binding pose, despite significant structural dynamics of the p7 bundle observed in hydrated lipid bilayers (Fig 3a). The root mean square deviation (RMSD) of the protein backbone Cα atoms indicted that the structural dynamics of JK3/32-bound p7 fluctuated within tolerable values, and were indistinguishable from unbound protein (data not shown). JK3/32 remained within the binding pocket throughout the course of the simulation, with the carbonyl group initially forming H-bonds with Tyr45 followed by subsequent bifurcation (after ∼50 ns) with Trp48 (Fig 3b). The JK3/32 amino group made further interactions with various side chains over the course of the simulation.

**Figure 3.**
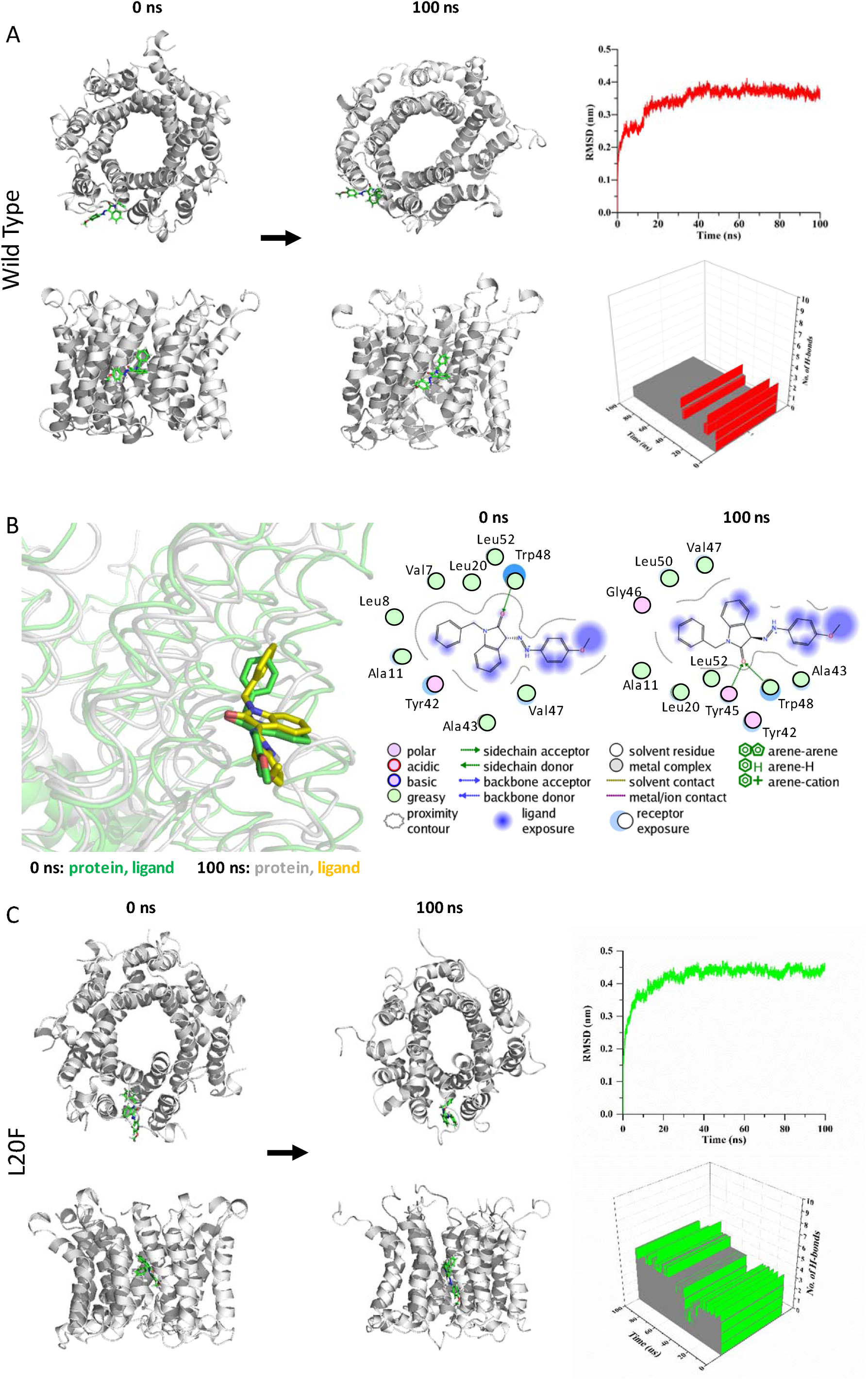
Atomistic simulations of JK3/32 interaction with PDB: 3ZD0-based heptamers. **A.** 100 ns atomistic molecular dynamics simulation of p7 complexes bound to JK3/32, starting from a minimised pose in a hydrated lipid bilayer environment. Top-down and side views of complexes at the start and end of simulation are shown with a single molecule of JK3/32 bound at the membrane-exposed binding site. Graphs on the right show the root mean-squared variation (RMSD) over time (top) and the number of H-bonds formed throughout the simulation. **B.** Overlay of p7 protein structures and JK3/32 orientation at 0 ns and 100 ns (left) alongside molecular interactions at the two time points (right); note bifurcation of H-bond interaction between indole carboxyl and Trp48 (0 ns) to both Trp48 and Tyr45 (100 ns). **C.** Effect of Leu20Phe rimantadine-resistance polymorphism upon JK3/32 interaction, set out as for A, above. Note, crowding of the binding pocket leads to higher stable RMSD and an increased number of H-bonds formed over time compared with the wild type protein (right panels).

Leu20Phe mutant p7 complexes also formed a stable interaction with JK3/32 (Fig 3c), although Phe20 caused reorganisation and crowding of the binding pocket stabilized by π-π stacking interactions between Phe20 and Tyr45. The intramolecular H-bond between the NH and carbonyl oxygen was lost on simulation, although bifurcated H-bonding to Tyr45 and Trp48 was maintained. We infer that efficient H-bond formation is contributory to JK3/32 potency. By contrast, p7 with the Leu20Phe mutation disrupted binding of rimantadine within the pocket, with the drug failing to make H-bond contacts and leaving the pocket over the course of the simulation (Fig S7).

### HCV entry is dependent upon p7 ion channel function

We previously demonstrated a link between p7 sequence and the acid stability of secreted HCV particles(*18*). This prompted us to speculate that, in addition to its role during secretion, virion-resident channel complexes might influence virus entry. In agreement, JK3/32, but not an inactive analogue from a closely related series, compound R21, reduced infectivity when added to infectious genotype 1b innoculae (Fig 4a, b). However, inhibition occurred at higher JK3/32 concentrations compared to those effective against virion secretion making it necessary to confirm effects were both HCV-as well as p7-specific. Reassuringly, the same genotype-dependent variation in JK3/32 potency was evident during entry experiments as observed during secretion; genotype 3a chimaeric viruses required 2-4 fold higher concentration than genotype 1b and R21 again had no effect (Fig 4c). As expected, compounds displayed no evidence of cytotoxicity at higher concentrations (Fig S8a, b). Note, JFH-1 was not employed during these experiments as R21 displayed modest but significant inhibitory activity against this strain in both secretory and entry assays, preventing its use as a negative control.

**Figure 4.**
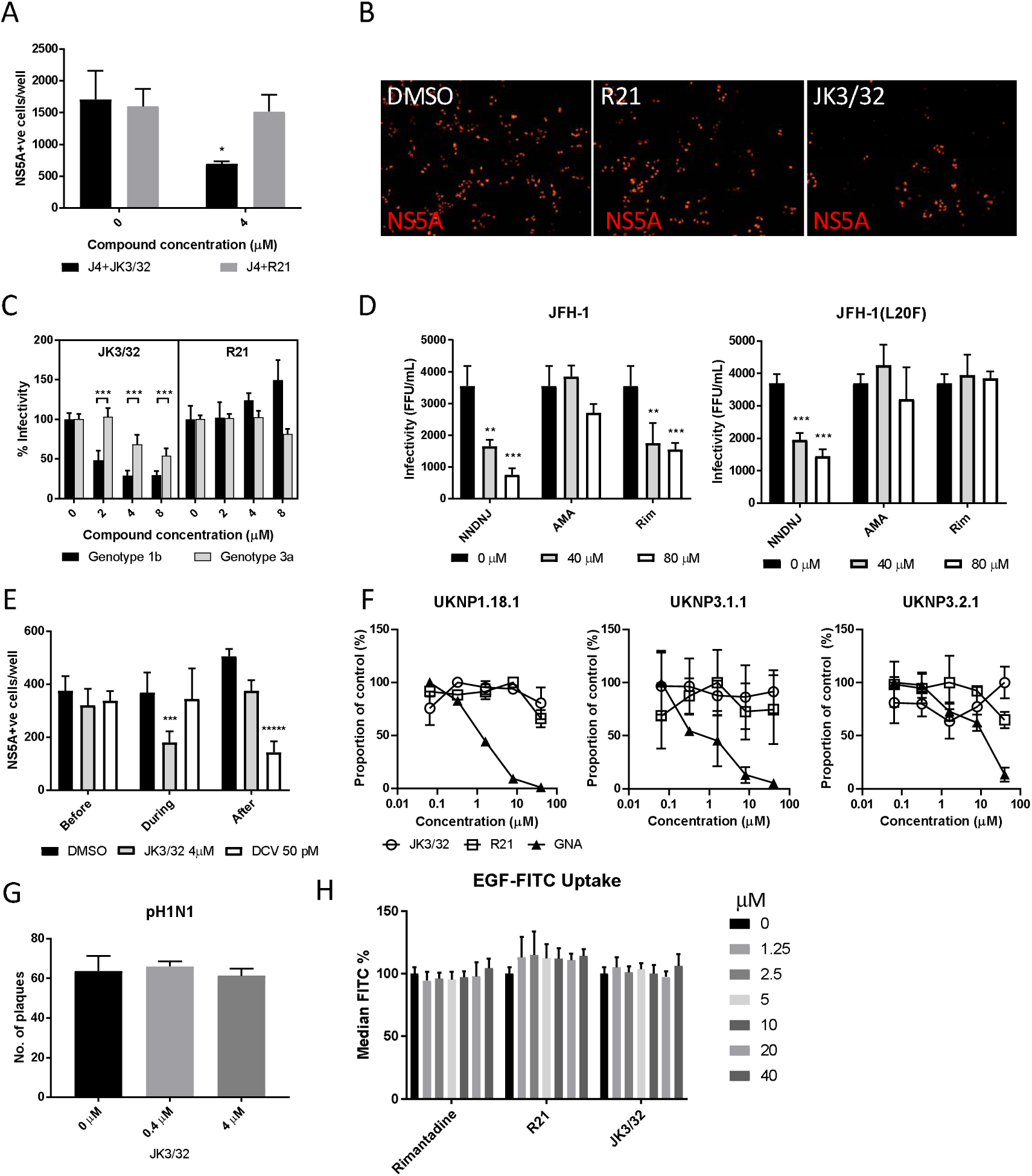
Characterisation of JK3/32 effects upon HCV entry. **A.** Infectivity following application of JK3/32, or the in ctive R21 analogue, during entry of GT1b chimaeric HCV into Huh7 cells. Virus innoculae were pre-treated with either compounds or DMSO for 20 min at room temperature prior to application to Huh7 cells overnight. Cells were washed extensively and assessed for infectivity 48 h post infection by NS5A immunofluorescence, quantified using the Incucyte Zoom. (*p≤0.05, Student T-Test, n=2). **B.** Representative images from Incucyte analysis of NS5A-stained Huh7 cells in A for comparison. **C.** Titrated JK3/32 concentrations were assessed during entry as described in A, comparing chimaeric GT1b (J4/JFH-1) and GT3a (S52/JFH-1) viruses (***p≤0.001, Student T-Test, n=2). **D.** Effects of prototypic p7 channel blockers against wild type and rimantadine resistant GT2a HCV (JFH-1) during entry into Huh7 cells (**p≤0.01, ***p≤0.001, Student T-Test, n=2). **E.** JK3/32 effects upon entry when added prior (2 h), during (4 h) or post-infection (48 h) with chimaeric GT1b HCV (J4/JFH-1), compared to Daclatasvir NS5Ai (***p≤0.001, *****p≤0.00001, Student T-Test, n=2). **F.** Lack of effect of JK3/32 or R21 upon entry of Lentiviral HCV pseudotypes into Huh7 cells compared to GNA positive control. HCV envelopes were derived from genotype 1b (UKNP1.18.1) or 3a (UKNP3.1.1 and UKNP3.2.1) patients. Results are representative of two experiments. **G.** Lack of effect for JK3/32 upon IAV entry into MDCK cells. **H.** Lack of effect for JK3/32, R21 or rimantadine upon clathrin-mediated endocytosis, measured by fluorescent EGF uptake.

Multiple unsuccessful attempts were made to select resistance to JK3/32 in culture (data not shown), meaning that it was not possible to provide genetic evidence supporting interactions with this compound during virus entry. We infer that this is due to the high degree of conservation seen within the JK3/32 binding site (Fig 2d). However, we previously defined both strain- and polymorphism-dependent resistance to prototypic adamantane and alkyl imino-sugar p7i(*23*), providing a means to assess p7 target engagement during entry. Secretion of genotype 2a JFH-1 HCV is innately amantadine-resistant, yet remains sensitive to rimantadine and the alkyl imino-sugar *N*N-DNJ. Virus entry experiments recapitulated this pattern, with the latter two compounds displaying dose-dependent efficacy (Fig 4d). Moreover, entry of JFH-1 Leu20Phe was resistant to both amantadine and rimantadine, whilst remaining *N*N-DNJ-sensitive; *N*N-DNJ disrupts p7 oligomerisation rather than binding peripherally(*23*). Hence, p7 sequence dictated p7i-mediated blockade of HCV entry, providing genetic evidence for virion-resident channels. Consistently, JK3/32 only blocked infection when introduced during virus infection, concordant with a direct effect upon virion-resident channels, whereas the NS5Ai Daclatasvir only interfered with HCV infection when added post-entry (Fig 4e).

Next, we used Lentiviral vectors pseudotyped with HCV glycoproteins to establish whether JK3/32 effects were p7-specific, or might instead interfere with receptor binding, particle integrity or membrane fusion. Encouragingly, neither JK3/32 nor R21 had effects upon the entry of pseudotypes possessing patient-derived E1/E2 representing genotype 1b or 3a(*38*), whereas *Galanthus nivalis* Agglutinin (GNA) blocked entry in a dose-dependent fashion (Fig 4f). Pseudotypes based upon prototypic genotype 1a or 2a, as well as the vesicular stomatitis virus glycoprotein (VSV-G) control, were also unaffected by JK3/32 or R21 (Fig S8c).

Lastly, to discern whether JK3/32 affected clathrin-mediated endocytosis, we tested sensitivity of pandemic H1N1 IAV entry to the compound. Not only does IAV enter cells via this pathway, it also retains virion-associated M2 viroporin proton channels thereby serving as an additional control for JK3/32 specificity. Accordingly, JK3/32 did not affect IAV entry (Fig 4g). In addition, JK3/32 did not interfere with clathrin-dependent uptake of fluorescent-tagged epidermal growth factor (EGF) (Fig 4h, S8d). Taken together, evidence supports that JK3/32 inhibits HCV entry in a p7 sequence-, genotype-, and temporally-dependent fashion, with no appreciable effects upon cellular endocytic pathways.

### JK3/32 and a labelled derivative exert irreversible blockade of virion-resident p7 channels

Despite multiple experiments supporting a specific effect upon HCV entry, the elevated concentrations of JK3/32 required to block infection made it desirable to minimise cellular exposure to the compound and so comprehensively rule out off-target effects. Fortunately, our in-depth understanding of JK3/32 SAR, docking, and MD simulations indicated that the creation of a labelled tool compound should be feasible. We predicted that the northern 4-OMe group should tolerate the addition of a flexible linker without significant loss to binding affinity. Click chemistry was used to attach an azide reactive fluorophore (Alexa Fluor 488 nm) to an alkynyl group at the 4-position of the phenyl ring generating a chemical probe (JK3/32-488), allowing compound concentrations to be calculated by fluorimetry.

Purified genotype 1b chimaeric HCV, dosed with 10 μM JK3/32, JK3/32-488, R21 or DMSO, was separated from unbound ligand by centrifugation through continuous iodixinol density gradients. Both control and R21-exposed gradients yielded overlapping peaks of infectivity at densities between 1.1 and 1.15 g/mL (corresponding to fractions 4-7 of the gradient), consistent with previous studies (Fig 5a, b, S9). However, infectivity was effectively abrogated in peak fractions treated with either JK3/32 or JK3/32-488, although a small number of infected cells were detectable (Fig 5a, b, S9). Quantification of the fluorescent signal from JK3/32-488 revealed concentrations within peak infectivity fractions were <2 nM (fractions 5 & 6). Hence, separation of treated virions from unbound compound using this method resulted in cellular exposure to far lower concentrations than those able to block HCV secretion, making off-target effects extremely unlikely (Fig 5c). Importantly, neither JK3/32 nor JK3/32-488 affected the stability of HCV particles as migration of virion-associated core protein corresponded with the peak of infectivity in each case, and was identical to the R21 control (Fig 5d). Hence, we conclude that effects upon HCV entry are not attributable to cellular exposure to JK3/32, and that the compound irreversibly blocks virion-resident p7 channels.

**Figure 5.**
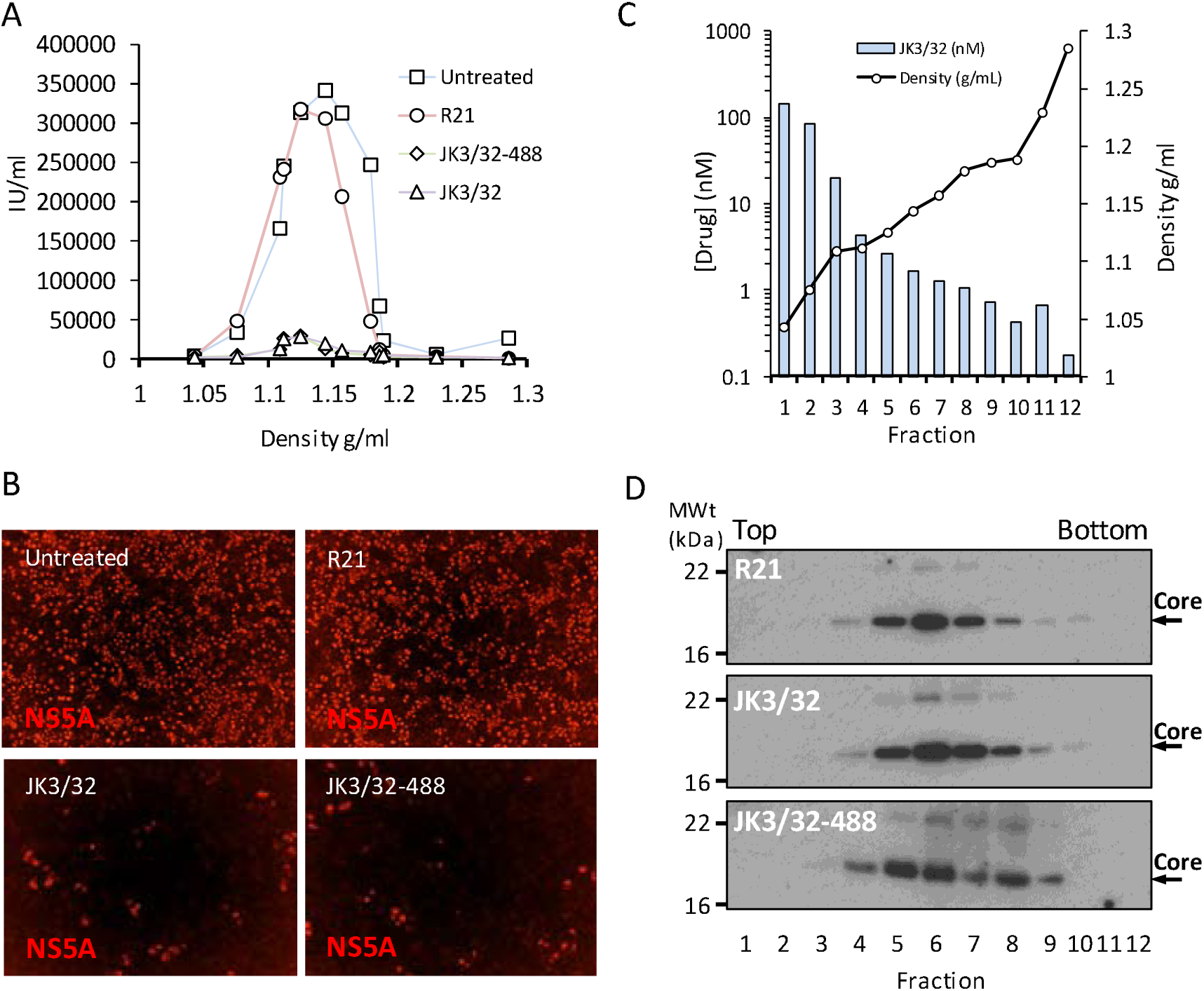
Direct JK3/32 virion dosing reduces HCV infectivity with minimal exposure of compound to cells. Concentrated, purified high titre chimaeric GT1b HCV (J4/JFH-1) was incubated with DMSO, active JK3/32(-488) or control compounds (R21) at 10 μM for 20 min prior to separation on a continuous 10-40% iodixinol/PBS density gradient followed by fractionation. **A.** IncuCyte quantitation of NS5A immunofluorescence staining within each fraction expressed as infectious units (IU) per mL. **B.** Immunofluorescence staining of HCV NS5A protein within naïve Huh7 cells 48 h post-infection with 10 μl of fraction 6, corresponding to peak infectivity for untreated control gradients. **C.** Calculated concentration of JK3/32-488 based upon fluorimetry within each of 12 fractions taken from the top of the gradient and corresponding fraction densities. **D.** Western blot analysis of drug treated gradient fractions, probing for HCV core protein.

## Discussion

Our rationally derived lead, JK3/32, and its associated SAR, represent a dramatic leap forward compared with previously described p7i on several fronts. First, correlation between SAR and the 3ZD0-based heptameric model supports hairpin-shaped protomers as the biologically relevant building blocks for channel complexes. Second, the pairing of potent p7i with chemically similar but inactive analogues provides well-controlled reagents with which to investigate the role of p7 channel function during the HCV life cycle; whilst previous hit compounds showed similar potency, they lacked a consensus pharmacophore(*24*). Third, SAR enabled creation of the first – to our knowledge – labelled tool compound based upon a viroporin inhibitor, further corroborating SAR and predicted 3ZD0 p7 interactions. Lastly, drugs based upon JK3/32 would represent a distinct antiviral modality compared with replicase-targeted HCV DAA, potentially targeting two discrete stages of the virus life cycle. Hence, this work sets a new precedent for viroporin-targeted drug development that should enable the field to advance beyond amantadine- and/or rimantadine-based influenza therapy.

The potency and selectivity demonstrated by JK3/32 was consistent with comprehensive SAR, docking predictions within the peripheral binding site, as well as MD simulations of the bound JK3/32 ligand over time. Each binding site occurs between adjacent hairpin-formation protomers and the majority of residues are highly conserved (Fig 2c, d). This includes Tyr45 and Trp48, which form key H-bond interactions with the JK3/32 indole carbonyl. Only Leu20 (∼46% conserved) shows appreciable variation within the binding site, with the rimantadine-resistant Phe20 occurring in just ∼3.6% of isolates in the EU HCV database (Fig 2d, Table S1); Val (∼30%) or Ala (∼12%) are more common. Moreover, Phe20 stabilises rather than disrupts interactions with JK3/32 during MD simulations, and JK3/32-related compounds are unaffected by this mutation in culture(*24*). We infer that the high degree of conservation within this site explains why JK3/32 specific resistance was not selectable in culture, despite several attempts.

Interestingly, the genotype 5a (EUH1480 isolate) 2M6X channel complex solution structure, comprising triple-helix protomers, contains Phe20 within a peripheral adamantane binding site(*29*); this site differs significantly in amino acid composition and structure compared to the 3ZD0-based hairpin heptamer model. Unfortunately, the recombinant protein used for structure determination did not display ion channel activity, making it impossible to determine drug sensitivity(*29*). Nevertheless, a related genotype 5a (SA-13) also retains Phe20 and is rimantadine-sensitive in culture(*37*), supporting that Phe20 mediated rimantadine resistance is context-specific. The EUH1480 p7 sequence was modified to enhance recombinant expression, primarily at Cys residues. Amongst these, Cys2 is only found within GT5a (present in just 0.07% of p7 sequences overall, see Table S1), and Cys27 is highly conserved across all genotypes (>95%). It is conceivable that these mutations were responsible for the lack of observable channel activity for this protein.

Unlike the peripheral rimantadine binding site in 3ZD0-based models, the region bound by amantadine in the 2M6X complex(*29*), comprising Phe20, Val25, Val26, Leu52, Val53, Leu55 and Leu56, is poorly conserved (Table S1). Only Leu52 and Leu55 represent consensus residues at their respective positions, with all the others being minority species. Moreover, three of the seven residues vary between the closely related SA-13 and EUH1480 genotype 5a strains (positions 26, 53 and 55). Unsurprisingly, JK3/32 binding and SAR is incompatible with this site within the 2M6X complex, or more importantly, with a genotype 1b homology model based upon the 2M6X structure (data not shown). Taken together, our findings add to the NMR-specific concerns recently raised regarding 2M6X(*30*) and support that biologically functional p7 channels comprise arrangements of hairpin protomers.

JK3/32 was compared to a patented p7i advanced into early phase human studies in South East Asia, BIT225(*36*) (N-Carbamimidoyl-5-(1-methyl-1H-pyrazol-4-yl)-2-naphthamide, Biotron Ltd, Australia). BIT225 is an amiloride-related compound selected as an inhibitor of genotype 1a p7 (H77 strain) from a bacterial screen, with activity against recombinant protein at 100 μM in suspended bilayers. As a surrogate for HCV in culture, BIT225 potently inhibited replication of the *Pestivirus*, bovine viral diarrhoea virus (BVDV) (IC_50_ 314 nM), although specificity was not attributed to the BVDV p7 protein(*36*). However, BIT225 later showed genotype-dependent efficacy versus HCV(*37*), with IC_50_ values of 10 μM or 30 μM (genotype 2a and 5a chimaeric HCV, respectively); interestingly, these concentrations are far higher than those effective versus BVDV(*36*). Whilst we observed a similar anti-HCV IC_50_ of 17.7 μM versus genotype 1b chimaeric HCV, this coincided with the CC_50_ of 18.6 μM (Fig S5), making it impossible to define a selective antiviral effect. Notably, Huh7 cell cytotoxicity was not determined in previous HCV studies(*37*), whereas BVDV studies reported a CC_50_ of 11.6 μM for Madin-Darby bovine kidney (MDBK) cells, with appreciable (∼16%) cytotoxicity at just 4 μM(*36*). We conclude that BIT225 may show greater potency versus BVDV than HCV, and that selective antiviral effects cannot be determined for HCV in the Huh7 system. This contrasts with the considerable selectivity index observed for JK3/32 (>500, Fig S2).

JK3/32 inhibition of secreted HCV infectivity showed cross-genotypic potency, including versus genotype 3a, which can be more difficult to treat using current DAAs (Fig 1b, c). Thus, we were naturally cautious regarding the higher concentrations required to block HCV entry, although this effect was reproducible compared to the inactive R21 analogue, showed genotype-dependent variation and was coincident with entry (Fig 4a-c, e). Lentiviral pseudotype assays supported that JK3/32 did not affect E1/E2 mediated receptor binding or membrane fusion (Fig 4f, S8c), plus it did not affect IAV entry or clathrin-mediated endocytosis (Fig 4g, h). Moreover, strain/polymorphism-dependent resistance to prototypic p7i provided genetic evidence of target engagement (Fig 4d) and we were effectively able to rule out cell-dependent effects using our fluorescent-tagged JK3/32 derivative (Fig 5), supporting irreversible dosing of virion-resident p7 channels.

We speculate that the different biological roles of p7 channel activity during secretion and entry may explain why higher p7i concentrations are required to block the latter process. During secretion, p7 must counteract active proton gradients generated by vATPase in order to prevent acidification within large secretory vesicles(*20*), conceivably requiring a considerable number of viroporin complexes. By contrast, during entry, virion-resident p7 would promote core acidification in the same direction as proton gradients generated within endosomes, presumably achieved by a small number of channel complexes. Thus, inhibiting a fraction of p7 channels within a secretory vesicle might prevent vATPase antagonism, whereas saturating inhibitor concentrations would be required during entry to prevent activation of even a single virion-resident p7 channel. This hypothesis may also explain why cell-to-cell infection appears less sensitive to p7i as virions presumably bypass acidifying secretory vesicles(*37*).

Studies of HCV harbouring mutations within the highly conserved p7 basic loop (consensus: K33, R35, p7 sequence) have provided evidence contrary to the presence of p7 channels within virions(*16*). Specifically, highly efficient chimaeric J6/JFH-1 harbouring basic loop mutations is able to produce secreted infectious virions, albeit at significantly reduced titre (∼ 4-log10 reduction). Basic loop mutant particles displayed equivalent specific infectivity compared with wild type, suggesting that p7 function is not required during entry. Moreover, in the less efficient JFH-1 background where basic loop mutants effectively abrogate secreted infectivity, restoration of a limited level of secreted infectivity occurred upon either *trans*-complementation with IAV M2, or Bafilomycin A (BafA) treatment of infected cells(*19, 20*). These observations suggest that supplementing p7 function during virion secretion is sufficient to restore infectivity, negating the requirement for p7 to act during entry.

However, this interpretation assumes that basic loop mutations specifically affect ion channel activity in isolation, yet no published evidence exists to support this notion. Instead, basic loop mutations induce defects in p7 stability and E2-p7-NS2 precursor processing within infected cells(*19*), as well as severe disruption of membrane insertion observed *in vitro*(*39*). It is likely that cell phenotypes directly link to defective membrane insertion, suggesting that p7 may spontaneously insert into membranes rather than depending upon the signal recognition particle. Spontaneous membrane insertion would explain why p7 lacking an upstream signal peptide forms functional channels within cells(*40*), as well as how HCV harbouring deletions of half of the p7 protein remain viable for replication despite the predicted effect of such mutations upon polyprotein topology(*41*). Thus, whilst only a minority of basic loop mutant p7 proteins might successfully insert into membranes, this should result in the formation of functional channel complexes. This scarcity of p7 complexes would profoundly disrupt virion secretion in the model proposed above due to an inability to counteract vATPase. However, a minority of virions might conceivably contain sufficient functional p7 complexes. Provision of M2 and/or BafA would therefore rescue secretion of such particles by allowing them to survive within acidifying vesicles, which could then proceed to infect naïve cells.

One limitation of our study is the absence of p7 immunological detection within purified HCV virions. We previously generated the only published p7-specific polyclonal antisera, which required extensive concentration to detect relatively high protein concentrations(*39*). Nevertheless, these are the only reagents described for the detection of native p7 protein, as confirmed by others(*8, 10*). Whilst we applied these reagents to virion detection, stocks expired prior to a conclusive outcome. Others used recombinant viruses expressing epitope tagged p7 to investigate the presence of channel complexes within virions by co-immunoprecipitation(*42*). However, whilst p7 was not detectable within either anti-ApoE or Anti-E2 precipitates, the levels of core protein present were also low; by analogy with M2, we expect p7 to be present in stoichiometrically low numbers compared with canonical virion components. Similarly, p7 was not detected by mass spectrometry (MS) conducted upon affinity-purified HCV particles(*43*), yet this was also true for the E1 glycoprotein, which should show equivalent abundance to E2 resulting from heterodimer formation.

Thus, poor reagent quality combined with low protein abundance hampers direct detection of p7 within virions, representing a significant challenge that we intend to address by the addition of an affinity tag to the “click”-labelled JK3/32 derivative. Concentration of virions and signal amplification will be essential as gradient infectivity peaks have an approximate particle molarity of 0.5 fM, based upon infectivity measurements (Figure 5a). IAV particles contain only 16-20 M2 proteins(*44*), meaning that even a high particle: infectivity ratio of 1000:1 would likely yield a maximal virion-associated p7 concentration in the tens of pM. Hence, a virion-associated fluorescence signal was not observed within peak infectivity fractions where the baseline unbound JK3/32-488 concentration was ∼2 nM.

The favourable potency and selectivity index demonstrated by JK3/32 could serve as a high quality starting point for a more comprehensive p7-targeted drug development programme. Blocking p7 activity would interfere with two distinct stages of the virus life cycle compared to existing replicase-targeted DAAs, namely virion secretion and entry. Hence, p7i could be ideal for delivering effective antiviral prophylaxis against *de novo* exposure (e.g. needle stick/iatrogenic or perinatal) or transplanted graft re-infection, in addition to use alongside conventional DAA combination therapies treating chronic infection. Whilst blocking both entry and secretion may require higher plasma concentrations compared with targeting secretion alone, the favourable selectivity index for JK3/32 suggests this should be feasible and future development may further optimise potency and bioavailability.

In summary, viroporins represent an untapped reservoir of antiviral drug targets that has historically been under-explored following the shortcomings of adamantane M2 inhibitors. Our work shows that it is possible to take a step-change in targeting these proteins, providing a new approach to antiviral therapy and generating novel research tools with which to dissect viroporin function.

## Materials and Methods

### Virus secretion inhibition assays

Huh7 cells were cultured and propagated as described previously(*21*). Secreted infectivity was measured as described(*34*). Briefly, 1 µg of linearised HCV constructs pJFH1, pJ4/JFH1, or pS52/JFH1 was used to perform *in vitro* transcription (RiboMax express, Promega) following the manufacturer’s protocol. Following phenol/ chloroform extraction, 4 × 10^6^ Huh7 cells were electroporated with 10 µg HCV RNA. Electroporated cells were seeded at 2.5 × 10^4^ cells/ well in 100 µL volume in 96 well plates and incubated 4 h. Compound dose responses were prepared at 400x in DMSO, diluted 1:20 into media in an intermediate plate, and 1:20 into the final cell plate to yield final 0.25% (v/v) DMSO. All compound treatments were dosed in duplicate. Dosed cells were incubated 72 h before performing 1:4 dilution (50 µL) of virus-containing supernatant onto a plate of naïve Huh7 cells (150 µL), seeded at 8 × 10^3^ cells/well 6 h previously. For cytotoxicity analysis, producer plates were washed in PBS and fixed in 4% PFA prior to imaging cellular confluency using an IncuCyte ZOOM (Essen BioSciences). Infected Huh7 cells were incubated 48 h before washing 3x in PBS and fixing in 4% PFA. Fixed cells were washed in PBS and permeabilised using 0.1% Triton X-100 (v/v) in PBS for 10 min, RT. Following PBS wash, cells were immuno-stained for NS5A to quantify infected cells. Anti-NS5A antibody was used at 1:2000 in PBS supplemented with 10% FBS, 16 h, 4°C. Following 3x PBS washes, AlexaFluor 594 nm Donkey anti-Sheep antibody was added at 1:500, 2 h, RT under subdued light. Cells were washed in PBS and imaged using phase and red channels (IncuCyte ZOOM). Infected cells positive for NS5A expression were quantified using IncuCyte parameters previously described(*34*), normalised to DMSO control and non-linear regression fitted using Prism 6 (GraphPad) to determine EC_50_/CC_50_. Each experiment included a serial dilution of untreated virus confirming 1:4 dilution fell within a linear range and internal DAA EC_50_ controls (data not shown). Determined Z-factor was routinely > 0.45. In addition, the Incucyte cell confluence tool was used as a measure of cell viability in both transfected/inhibitor-treated producer cells and target cell populations; producer cell plates were also subjected to MTT (3-(4,5-dimethylthiazol-2-yl)-2,5-diphenyltetrazolium bromide) assays to determine potential compound effects against Huh7 cell metabolism. Lastly, analogous plates were set up using Huh7 cells electroporated with a subgenomic HCV replicon (genotype 2a JFH-1 strain – the same NS3-5B replicase as present within chimaeric HCV) encoding firefly luciferase in the first open reading frame. 48 h post-electroporation, cells were lysed and luciferase activity determined using a commercially available kit according to the manufacturer’s instructions (Promega).

### HCV entry inhibition assays

HCV entry experiments used virus supernatant stocks harvested 72 hours post-electroporation and stored at -80°C. Huh7 cells were seeded at 8 × 10^3^ cells/ well in 100 µL in 96-well plates for 6 h. Indicated compound was added to virus-containing supernatant stocks immediately prior to infection at MOI of 0.6 and incubated 18 h. For “Time of Addition” experiments (Figure 4e), inhibitors were added for 2 h *before*, for 4 h *during*, and for 48 h *after* infection, as indicated. Cells were washed and incubated 48 h in media prior to quantifying infected cells via NS5A immunostaining as described above. Statistical significance was determined for test samples vs. controls using unpaired, two-tail Student T-tests. Note, incubation for 48 h was necessary to achieve robust and reproducible numbers of infected cells for counting in IncuCyte assays; shorter assays led to unacceptable errors in quantitation. However, as shown in Figures S1 and S2, the assay was proven to discriminate p7-dependent effects from those resulting from impaired RNA replication or cell viability.

### Liposome dye release assay

Recombinant genotype 1b p7 (J4 strain) was expressed and purified as described previously(*11*). Channel activity was assessed using liposome dye release assays(*21, 35, 39*), mixing protein with liposomes containing a self-quenching concentration of carboxyfluorescein and monitoring ensuing gain in fluorescence as an indirect measure of p7 activity.

### Influenza A virus Plaque reduction assay

Madin-Darby Canine Kidney (MDCK, from ATCC) cells (seeded at 5 x10^5^ / well of a 12 well plate 4 hours prior to infection) were infected for 1 h with A/England/195/2009 (E195) influenza A virus (IAV) at a multiplicity of infection (MOI) of 0.01, following preincubation with compounds for 30 min on ice. Virus containing media was then removed and replaced with serum free (SF) minimal essential media (MEM) with 1 µgml^-1^ TPCK trypsin containing compounds for 24 h. Clarified supernatant dilutions of 10^-1^ to 10^-4^ were then used to infect fresh monolayers of MDCK for 1 h, then replaced with a 3:7 mixture of 2% w/v agar (Oxoid™ Purified Agar) and overlay media (MEM, 0.3 % v/v BSA (fraction V), 2.8 nM L-Glutamine, 0.2 % v/v NaHCO2, 14 mM HEPES, 0.001 % v/v Dextran, 0.1x Penicillin and 0.1x Streptomycin) containing 2 µgml-1 TPCK trypsin. Agar was removed after 72 h and cells fixed in 2 ml 4 % paraformaldehyde in PBS for 1 h prior to staining with 1 % v/v crystal violet solution for 5 min, enabling plaques to be visualised and counted.

### Lentiviral pseudotype entry assays

Murine leukaemia virus (MLV) based Lentiviral pseudotypes were generated using a three plasmid system comprising a luciferase reporter vector (pTG126), an MLV Gag-Pol expressing plasmid (phCMV-5349) and a plasmid encoding the vesicular stomatitis virus glycoprotein (VSV-G) as a positive control, relevant HCV E1/E2 sequences (pcDNA3.1D-E1/E2), or lacking an envelope as a negative control. Two mg of each plasmid were mixed and transfected using polyethylenimine (PEI) into HEK293T cells, seeded the previous day in a 10 cm culture dish (1.2 × 10^6^ per dish). Transfections were removed after 6 h and media replaced (Dulbecco’s Modified Eagle Medium with 10 % v/v foetal calf serum and non-essential amino acids). Supernatants were harvested at 72 hr post-transfection and clarified through a 0.45 μm filter prior to use.

Pseudotypes were generated using VSV-G, HCV envelopes from prototypic strains (genotype 1a (H77) and genotype 2a (JFH-1 and J6)), as well as three patient isolates UKNP1.18.1 (genotype 1b), UKNP3.2.1 and UKNP3.1.1 (both genotype 3a) as described previously(*45*). Pseudotypes were treated with the indicated final concentrations of inhibitor (JK3/32, R21 or GNA) for 90 min prior to transduction of Huh7 cells in DMEM for 4 h at 37 °C. Following transduction, the media/pseudotype/inhibitor mix was removed and cells incubated in fresh DMEM for 72 h before lysing and measuring luciferase activity in the HCVpp treated cells. Assays were normalised to positive controls lacking inhibitor and baseline determined using a pseudotype preparation with no E1/E2 present (delta-E). All conditions were performed in triplicate and data shown are representative of two independent experiments.

### Virus purification and ultracentrifugation

High titre J4/JFH-1 virus stocks were generated by sequential daily harvest of Huh7 culture supernatants over a 1-week period, comprising 20 mL of HEPES-buffered media in each of eight T150 cell culture flasks, seeded with 2 × 10^6^ electroporated cells on day 1. Supernatants were clarified prior to addition of 1/3^rd^ volume of 30% (w/v) polyethylene glycol (PEG)-8000 in PBS, thorough mixing and incubation at 4°C overnight. The next day, precipitates were spun at 2000 rpm for 40 min at 4°C in a Hereas bench-top laboratory centrifuge, pellets harvested, and then resuspended in 1/100^th^ the original culture volume of PBS. Stocks were titred using the IncuCyte and snap-frozen in dry ice/ethanol prior to storage at -80°C.

Approximately 2×10^6^ IU of virus were diluted into 200 μL PBS and treated with either DMSO or p7i at a final concentration of 10 μM. Suspensions were then layered over a pre-formed 10-40 % (v/v) iodixinol gradient in a 2.2 mL open-topped mini-ultracentrifuge tube. Gradients were then centrifuged at 150 000 x *g* for 3 h at 4°C in a S55S Sorvall rotor prior to harvesting into twelve equal fractions. 10 μL was removed for infectivity testing and 50 μL for fluorimetry after adjusting to 0.1 % (v/v) Triton-X100 to lyse virions. Blank gradients run in parallel were fractionated as above, then 100 μL per fraction was placed in labelled pre-weighed 1.5 mL Eppendorf tubes and resultant mass determined for density calculation. The remainder of gradient fractions were mixed with an equal volume of 2 x Laemmli SDS-PAGE sample buffer and stored at -20°C prior to thawing and analysis of 10 μL/fraction by SDS-PAGE and western blotting.

### SDS-PAGE and western blotting

Gradient samples were separated on a 4-20% Tris-Glycine acrylamide gel using a BioRad MiniProtean III rig, at a set voltage of 120 V for 60-90 min. Gels were then dismantled and placed within a pre-wetted sandwich of thick blotting paper on top of a pre-cut PVDF membrane (0.45 μm) that had been activated in 100 % MeOH for 5 min at RT. Proteins were transferred for 2 h using a Hoeffer semi-dry transfer rig set at 320 mA. Blotting sandwiches, gels and membranes were thoroughly pre-soaked in transfer buffer (1 x Tris-Glycine pH 8.3, 20 % MeOH) prior to assembly. Membranes were removed from transfers, and placed in 20 mL blocking solution (5 % w/v fat-free milk in 1 x Tris-buffered Saline, 0.1 % Tween 20 (TBS-T)) and shaken at RT for at least 3-4 h. Membranes were then washed in TBS-T and placed in 10 mL of blocking solution containing primary antibody (mouse anti-HCV core, C7-50, Thermo Fisher catalogue # MA1-080) diluted 1/1000, at 4°C overnight with gentle shaking. The next day, primary antibody was removed by three washes in 1 x TBS-T at RT for 10 min, followed by incubation with secondary antibody diluted 1/5000 in blocking solution (goat anti-mouse HRP conjugate, SIGMA) for 2 h at RT. Washing was then repeated prior to visualisation using ECL prime chemiluminescence substrate (GE Healthcare/Amersham), according to the manufacturer’s instructions.

### EGF uptake assay

Huh7 cells were seeded into 24-well plates (1.5×10^5^ per well) and left to adhere for ∼6 h. Cells were then treated using titrated compounds or Bafilomycin A (1μM) as a positive control for 4 h. Cells were then washed extensively and media replaced with PBS containing 2 μgμL^-1^ FITC-conjugated epidermal growth factor (EGF, Invitrogen) for 30 min. Cells were then washed extensively, removed using a cell scraper and then fixed in 4 % (w/v) paraformaldehyde in PBS for 10 min at room temperature. Flow cytometry was then used to determine median FITC levels, with gating on intact cells. All conditions were performed in triplicate.

### Commercial kinase activity screen

JK3/32 was tested at the MRC Protein Phosphorylation Unit International Centre for Kinase Profiling, Dundee, Scotland (http://www.kinase-screen.mrc.ac.uk/services/express-screen). The express screen was undertaken (50 human kinases) using 10 μM JK3/32 in a highly accurate radioactive filter binding assay using ^33^P ATP. This method is sensitive, accurate and provides a direct measure of activity. A control plate using reference compounds is analysed in parallel for quality control purposes. Values for % kinase activity are then determined.

### Analysis of p7 sequence conservation

1456 aligned HCV p7 sequences from all genotypes were obtained from the EU HCV database website in FASTA format. Sequences were aligned and visualised using Jalview (www.jalview.org), allowing the percentage occupancy at each of the 63 amino acid positions within the protein to be quantified (Table S1, Fig S6). Data was exported into MS Excel, then inputted into a free sequence logo website (https://weblogo.berkeley.edu/) to visualise relative occupancy (Fig 2d).

### Prototypic p7i

Rimantadine hydrochloride was purchased from Chembridge, amantadine hydrochloride from SIGMA and *N*N-DNJ from Toronto Biochemicals. BIT225 was synthesised via 5-bromo-2-naphthoic acid and 5-(1-methyl-1H-pyrazol-4-yl)-2-naphthoic acid according to the patent detail (US20150023921A1); analytical data was consistent with the expected structure.

### Generation of optimised heptameric 3ZD0-based p7 channel structure

The initial heptameric channel model was constructed using the Maestro programme (Schrödinger) based upon the monomeric unit from the 3ZD0 NMR structure as described(*24*). A heptameric bundle arrangement of protomers was oriented with His17 oriented towards the lumen equidistant from a centroid atom placed in the middle of the lumen to serve as a rotational centre with multiple energy minimisation steps. Iterative rotation of protomers in an octanol environment generated solutions. The preferred model was then refined using molecular dynamics simulations (methods described below).

### Structure guided molecular dynamics simulations and compound docking

#### Molecular Docking

The p7 structure was energy minimized using the default energy minimization scheme in MOE software (Version 2015:1001, www.chemcomp.com), AMBER 94 force field was used with □ = 2. Hydrogen atoms were added and partial charges were assigned using MOE. The energy minimization was carried out by restraining protein backbone atoms as rigid. The pairwise alignment of the structures, the one before minimization and the one after minimization, gives an RMSD of 0.10 Å. Ligands were prepared using MOE software. Hydrogens atoms were added using the protonate 3D option and then partial charges were assigned with MMFF94 force field default parameters. Energy minimizations were performed using MMFF94 force field with a root mean square gradient of 0.1 kcal Mol^-1^.Å^-1^ and gas phase solvation model. For docking, Leu20 or Phe20 residues were selected from each monomer of the wild type and mutated proteins respectively. Receptor sites with a radius of 5 Å is defined around Leu20 and Phe20 residue of each chain. The energy-minimized ligands were loaded into the MOE graphical user interface. The ‘rigid receptor’ protocol of MOE was used, where side chains of the protein were kept ‘fixed’ during the force field base refinement. Ligand placement was performed using the Triangle Matcher protocol, where the active site was defined using the α-spheres and the centre of that spheres are triplets of atoms. These triplets represent the location of tight packing. Poses are generated by superposing the ligand atom triplets onto the centre of the α-spheres. The top 1000 poses received after the placement were then used to score using the London dG scoring function which is an energy term summing contributions from ligand entropy from its topology or ligand flexibility, geometric imperfections of protein-ligand and metal-ligand interactions and desolvation energy. The top 30 poses ranked accordingly based on the London dG scores are then taken for a forcefield based refinement within the rigid receptor. The resulting poses are then rescored using Generalized Born Volume Integral/Weighted Surface Area dG (GBVI/WSA dG) scoring function. At the end final 30 poses were ranked accordingly while removing the duplicate poses.

#### MD simulations

To simulate all the systems, Amber ff99SB-ILDN force field (ff) with Amber/Berger combination of ff was used with Gromacs 4.5.5. The POPC lipid ((1-palmitoyl-2-oleoyl-sn-glycero-3-phosphocholine) topology was received from Cordomí *et al.*(*46*) while for the calculation of ligand parameters the generalized amber force field (GAFF) was used. The partial charges (ESP) were estimated using a HF/6-31G* basis set and Antechamber package was used for restrained electrostatic potential (RESP) fitting. Each of the bundles, (wild type docked with JK3/32 and Rimantadine, L20F mutated bundle docked with JK3/32 and Rimantadine, and only protein) was inserted into patches of hydrated and pre-equilibrated POPC lipids. The overlapped lipids were removed and after steps of minimization (2000 steps of steepest decent and 5000 steps of conjugated gradient), it was equilibrated for a total of 1.9 ns by restraining the positions of the heavy atoms proteins and ligands. The systems were brought to equilibrium by gradually increasing the temperature of the systems from 100 K to 200 K and 310 K. During the initial equilibration, the peptides and ligands were fully restrained by applying a harmonic potential with a force constant, k = 1000 kJ mol^-1^ nm^-2^. The simulations at temperatures 100 K and 200 K were run for 200 ps each followed by simulation at 310 K for 500ps. Once the systems were equilibrated at 310K, the restraints were released by running 3 short (500 ps) simulations, one with k = 500 kJ mol^-1^ nm^-2^ the consecutive two simulations with k = 250 kJmol^-1^ nm^-2^ and finally 0 kJmol^-1^ nm^-2^.

For all the simulations SPC/E water model was employed and ion parameters proposed by Joung *et al.* were adopted(*47*). A Nosé-Hoover thermostat with a coupling time of 0.1 ps coupling separately to the temperature of the peptide, lipid, and the water molecules and Berendsen barostat with a coupling time of 2.0 ps were used during the MD simulations. For the systems without any docked ligand and with JK3/32 docked, 14 Cl-ions were added to neutralize the simulation box while for the systems with rimantadine docked it was 15 Cl-ions. All the systems were consisted of 449 lipids which were hydrated with 14605-14606 water molecules (with 14 and 15 Cl-ions respectively).

### Compound synthesis and purification

Reagents and solvents were obtained from commercial suppliers and used without further purification. Thin layer chromatography (TLC) analyses were conducted using silica gel (aluminium foil backing) plates and visualised under UV radiation in a dark-box. Compound purification was effected using gradient elution on a Biotage Isolera-4 running SiO_2_ cartridges. HPLC-MS was performed on a Bruker Daltronics spectrometer running a gradient of increasing acetonitrile (0 to 100%) in water containing 0.1% TFA at 1 ml.min^-1^ on a short path ^18^C reverse phase column detecting compounds with both a diode array detector and a Bruker Mass spectrum analyser. HRMS experiments were conducted on a Bruker MaxisImpact time-of-flight spectrometer operating in a positive ion mode with sodium formate as an internal standard. ^1^H, ^13^C experiments were recorded using either a Bruker DRX 500 instrument or a Bruker DPX 300 operating at 298K, by solubilising the sample in deuterated chloroform (CDCl_3_) with internal standard tetramethylsilane (TMS), CD_3_OD, d_6_-DMSO, or d_6_-acetone as the NMR solvent. Chemical shifts were expressed as parts per million (ppm) and the splitting signals of NMR assigned as s (singlet), d (doublet), t (triplet), dd (doublet of doublet), br (broad) or m (multiplet).

## Acknowledgments

**General**: We are grateful to Jens Bukh (Hvidovre University Hospital and University of Copenhagen, Hvidovre, Copenhagen, Denmark), Takaji Wakita (National Institute for Infectious Diseases, Tokyo, Japan) and Wendy Barclay (Imperial College, London) for the generous provision of reagents. We thank Adrian Whitehouse (Leeds) for useful discussion. We also thank Morgan Herod and Adam Davidson (Leeds) for technical advice regarding the Incucyte Zoom.

## Author contributions

Performed experiments/simulations: JS, RG, MMK, JK, DRM, TLF, CS, BJK, EB, MJB, LW, AB, AWT, SG Interpreted and analysed data: JS, AWT, WF, RF, SG. Experimental concepts and planning: SG, RF, WF, JM, AWT, AM Reviewed manuscript: AS, MH, DR, AM, AWT Wrote manuscript: SG, RF, WF

## Data and materials availability

In-house synthesised compounds, where stock is available, are obtainable via MTA.

## Supplementary Figures

**Supplementary figure 1.**
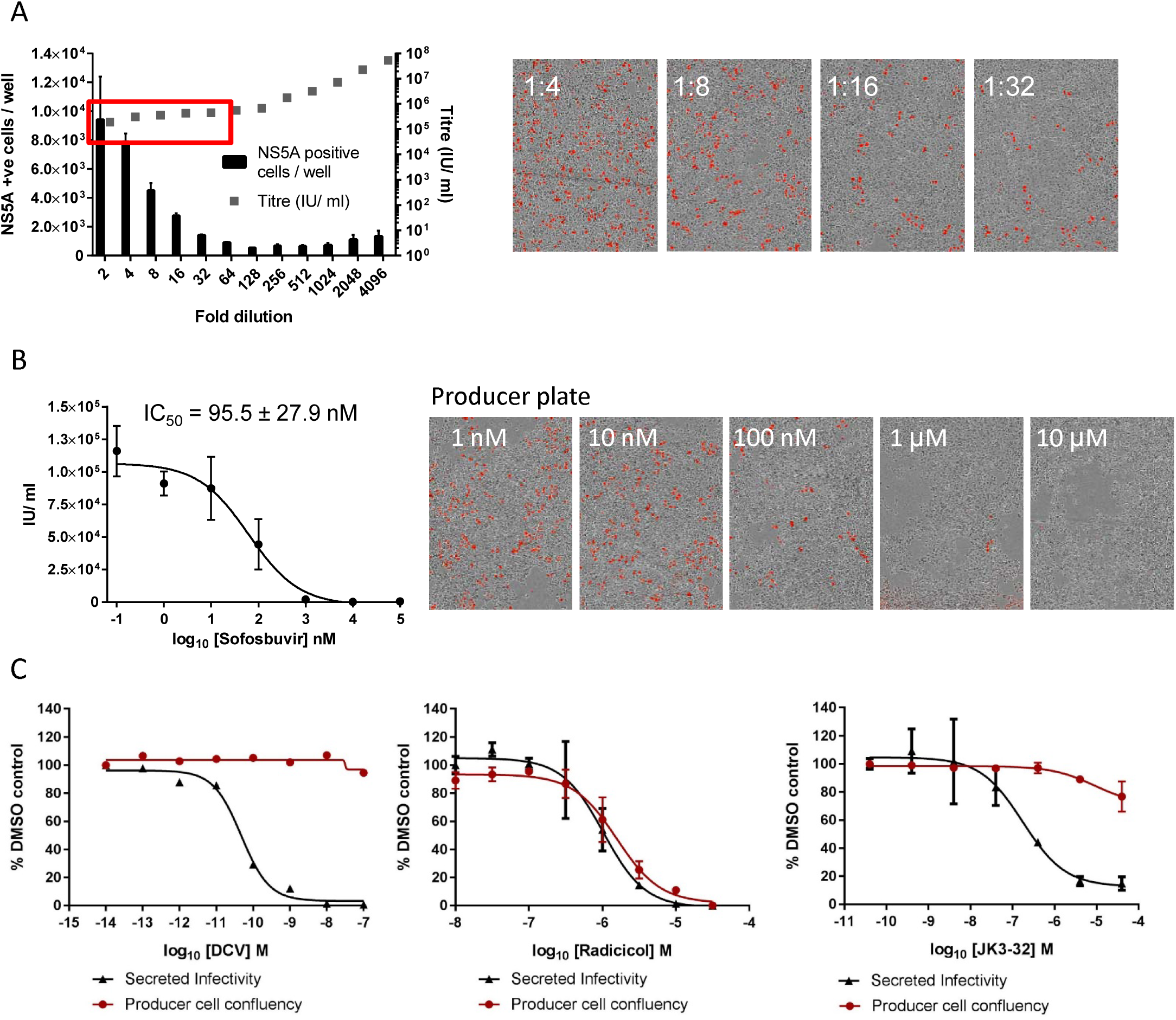

**Supplementary figure 2.**
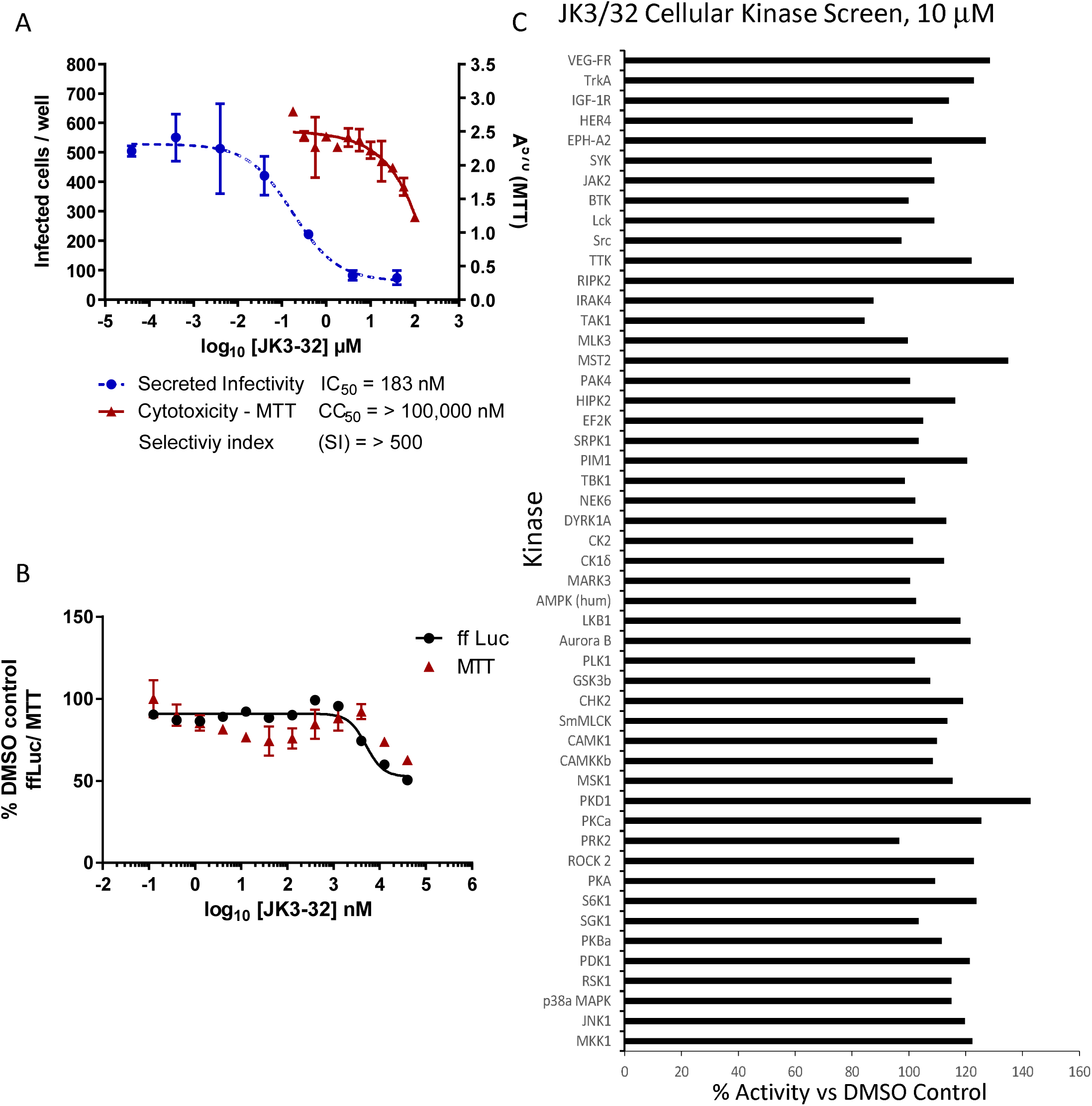

**Supplementary figure 3.**
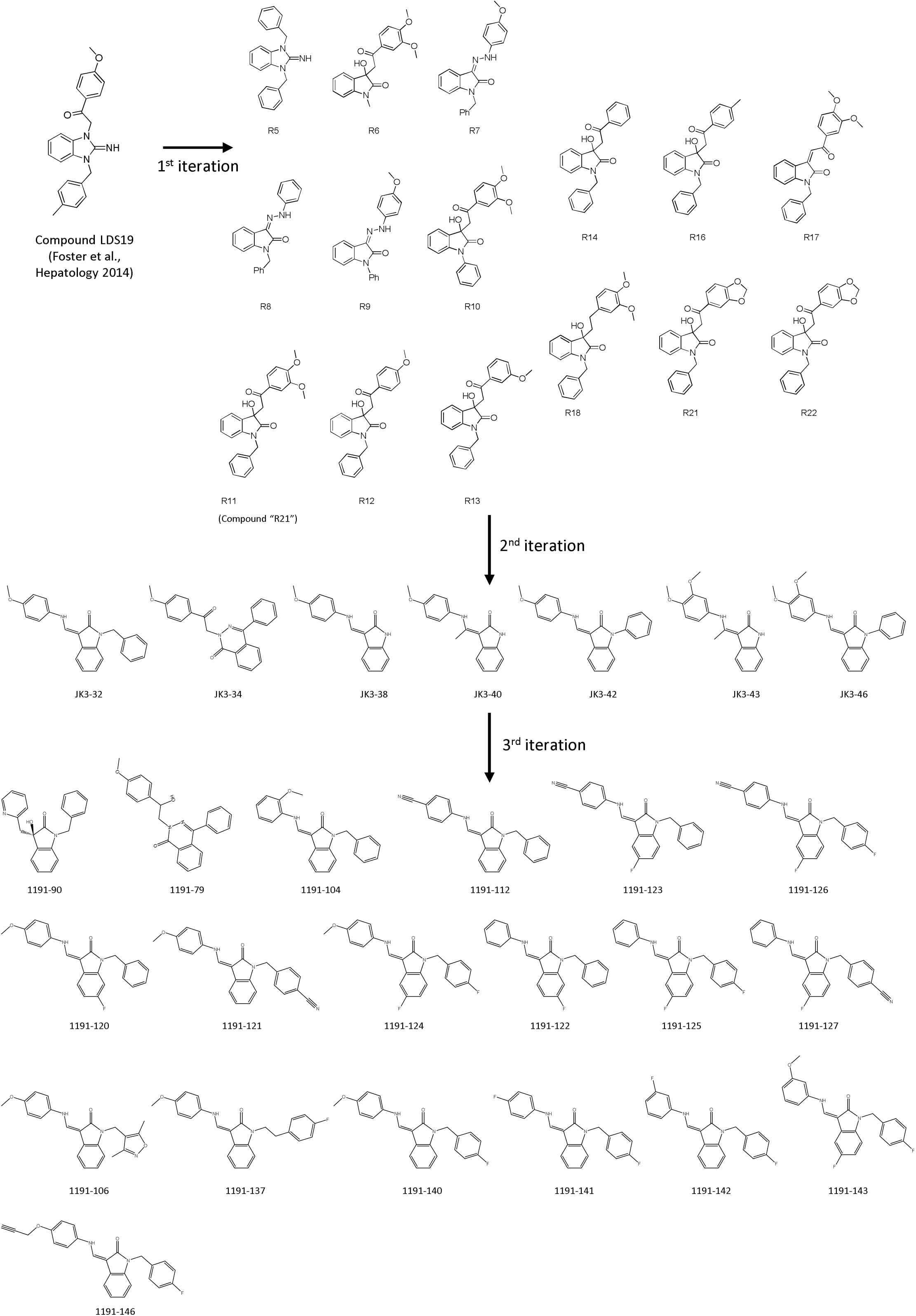

**Supplementary figure 4.**
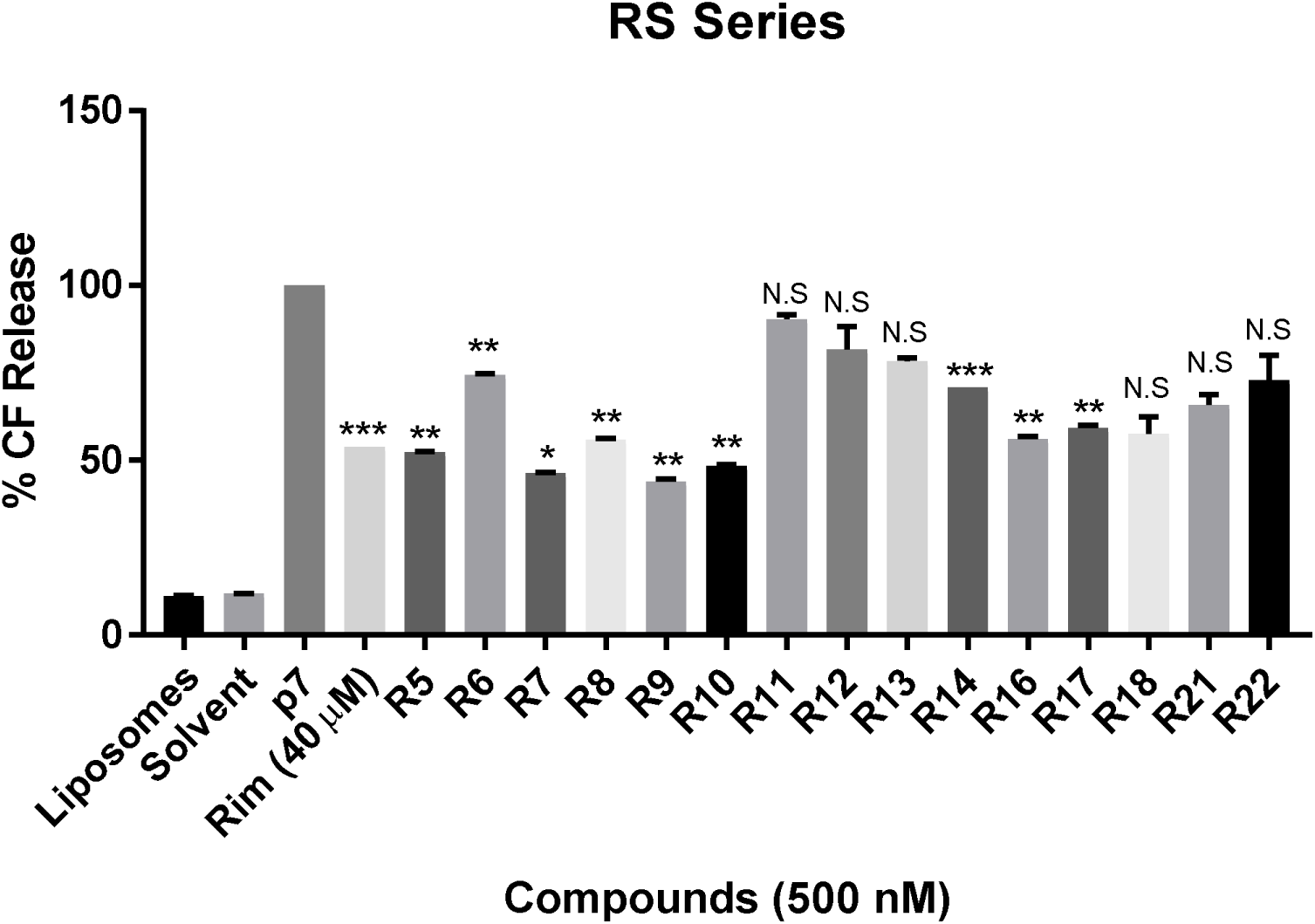

**Supplementary figure 5.**
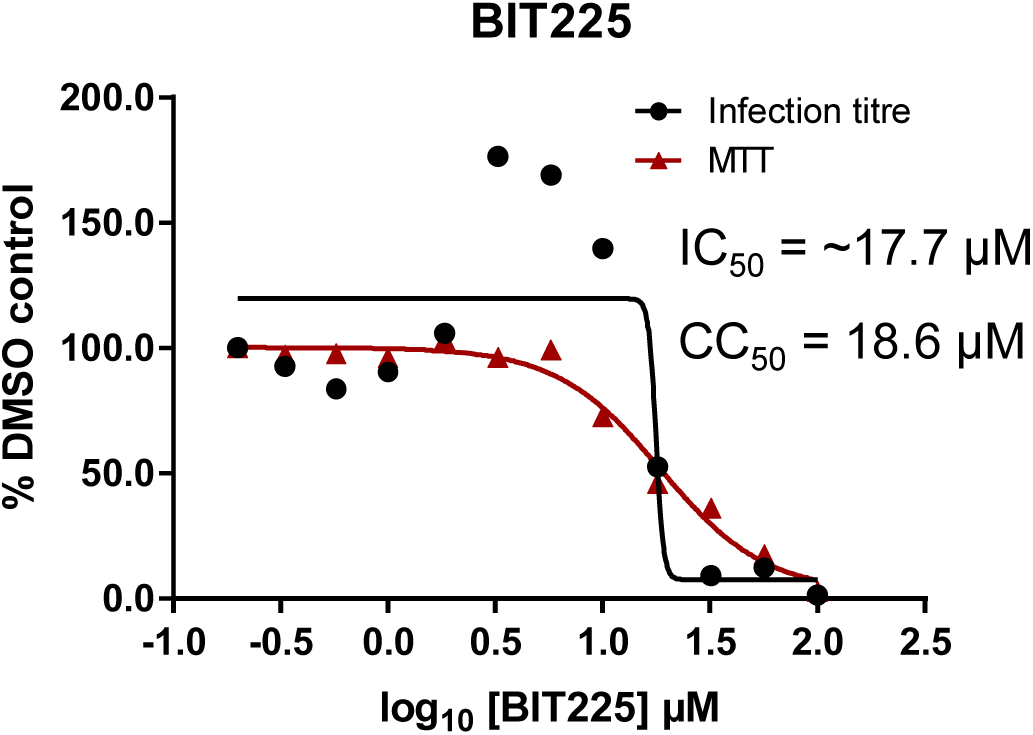

**Supplementary figure 6.**
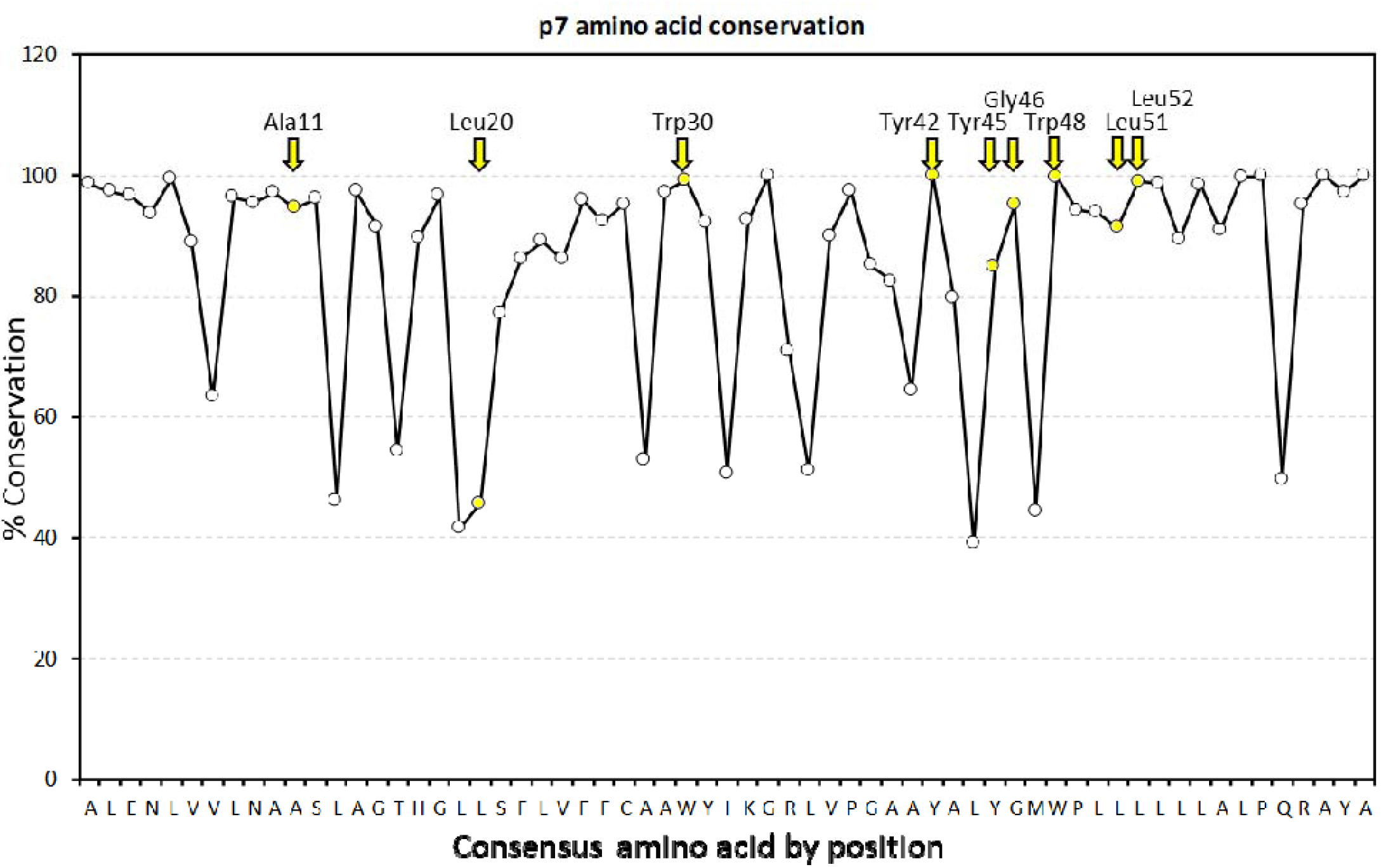

**Supplementary figure 7.**
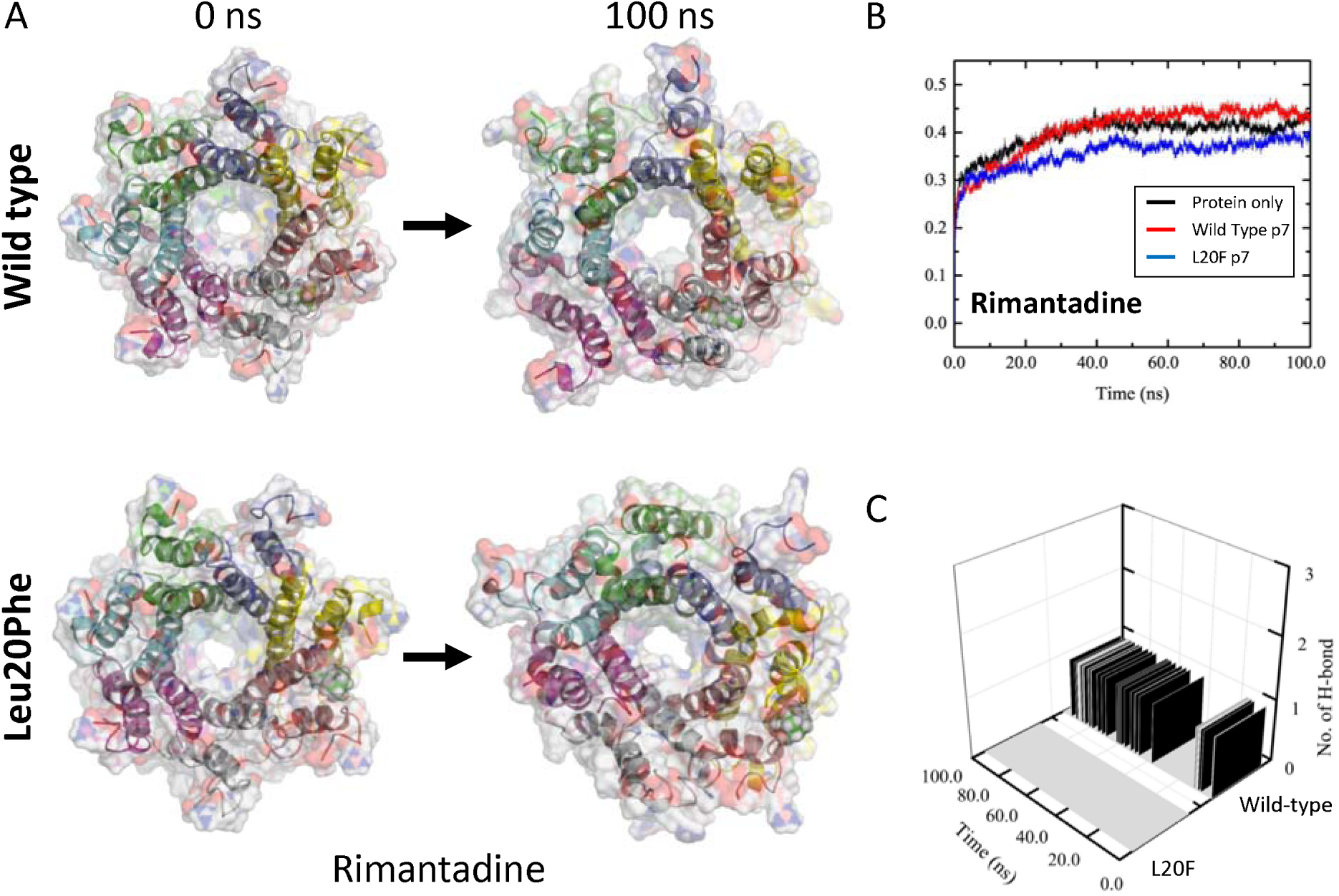

**Supplementary figure 8.**
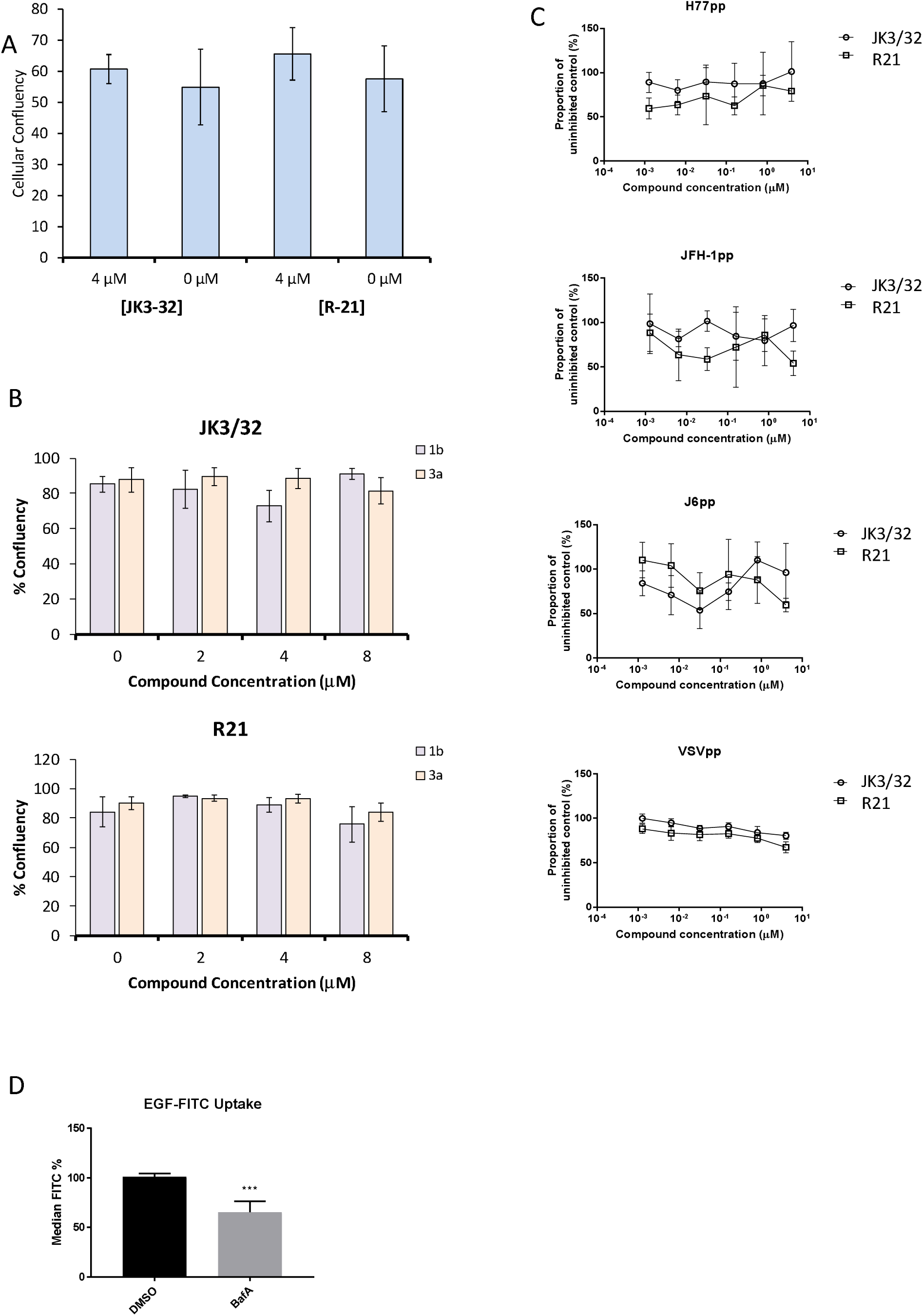

**Supplementary figure 9.**
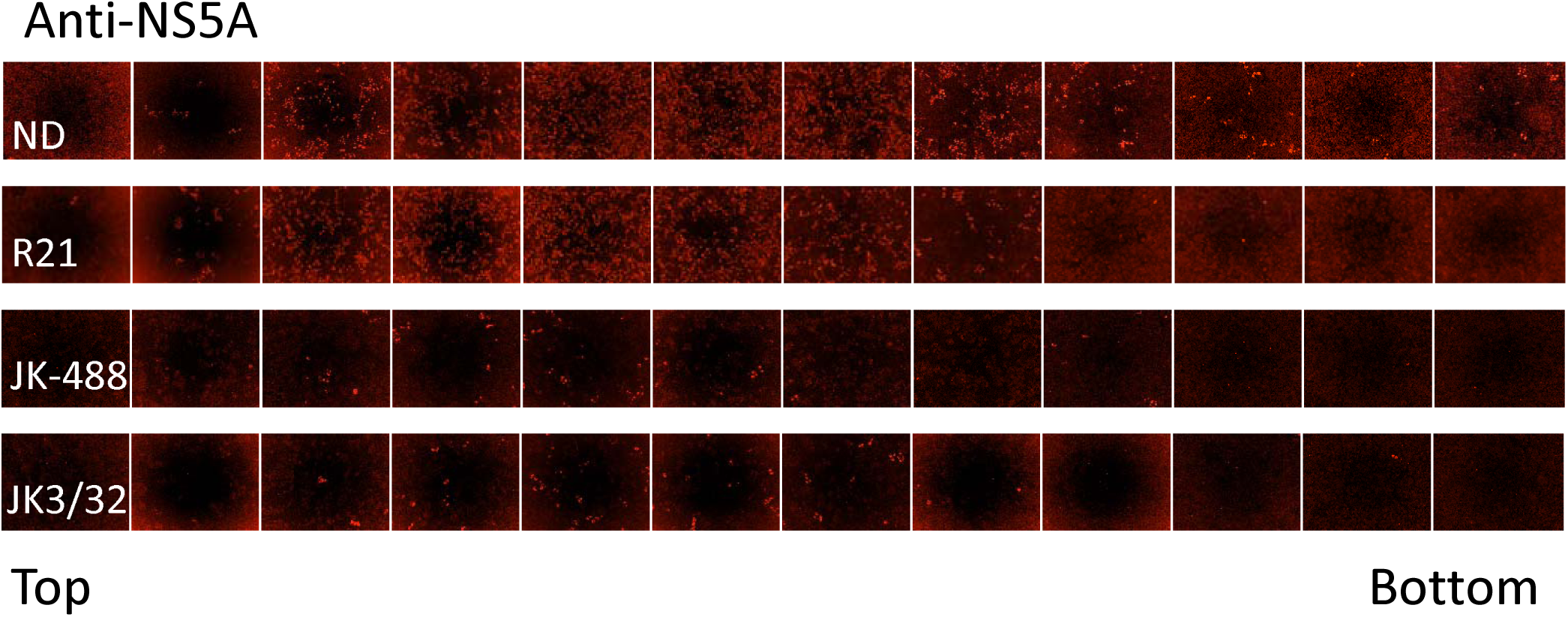

