## Supplementary figures for "Rationally derived inhibitors of hepatitis C virus (HCV) p7 channel activity reveal prospect for bimodal antiviral therapy"

**Supplementary Material**

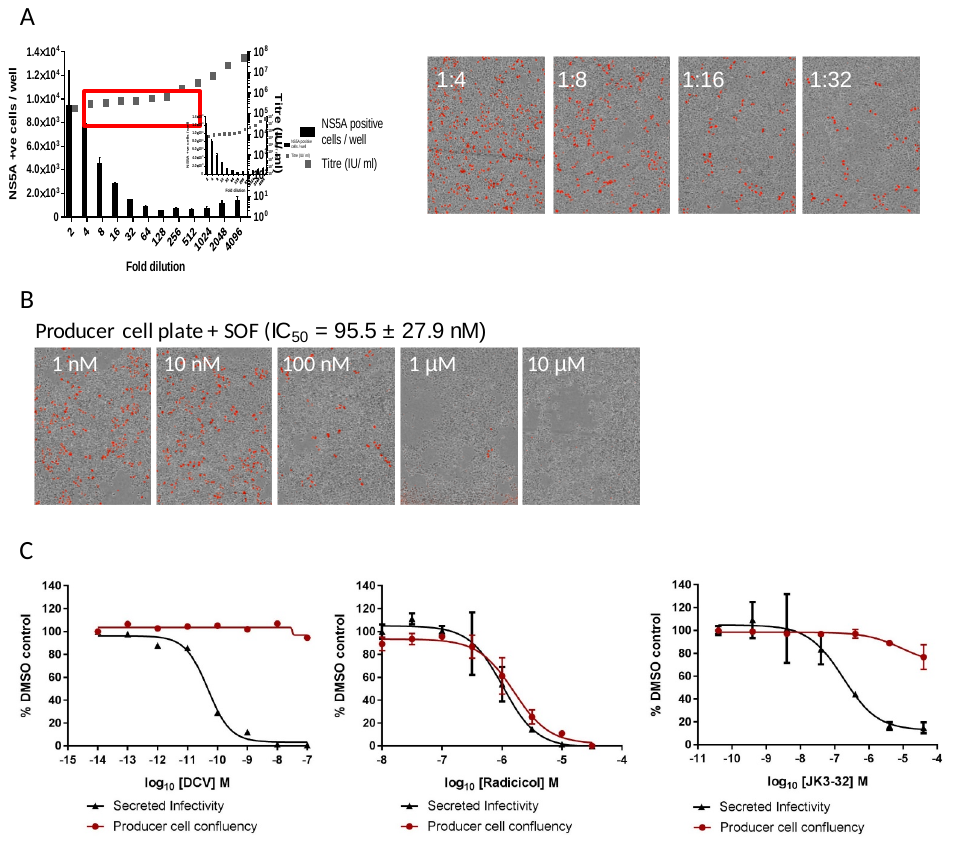

**Figure S1.** *Optimised quantitation of secreted HCV infectivity using the IncuCyte Zoom*. **A.** Dilutions of infectious supernatants were assessed for optimal signal/noise whilst remaining within the linear range of quantitation (red box). Appropriate dilutions for untreated virus were repeated alongside every inhibitor IC_50_ experiment to ensure that titres could be back-calculated accurately; each condition repeated in quadruplicate across multiple experiments, error bars show standard deviation. Example IncuCyte images are shown for comparison. **B.** IncuCyte images of Sofosbuvir (SOF) treated producer cells stained for NS5A, correlating infectivity with producer cell replication. **C.** Determination of cytotoxic effects by producer cell confluency. In each case, infectivity (black line) was assessed alongside producer cell confluency (red line) as an indicator of viability (MTT assays were also conducted for compounds showing effects in initial screens). Examples shown for a control DAA Daclatasvir (DCV), the Hsp90 inhibitor Radicicol as a Huh7 cytotoxic agent, and the lead p7i JK3-32. Error bars show standard deviation for quadruplicate samples in each representative experiment.

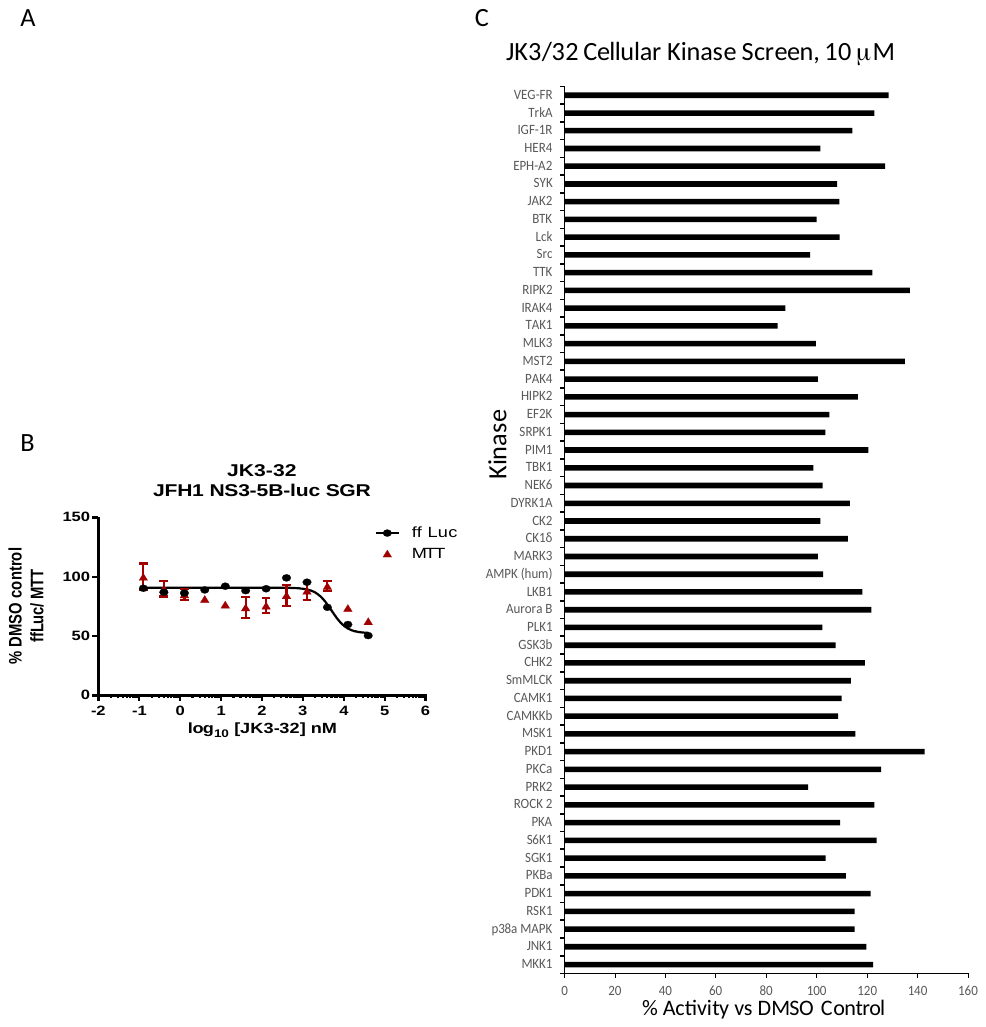

**Figure S2.** *Screening for potential JK3/32 off-target effects*. A. Summary data establishing the JK3/32 selectivity index with respect to cytotoxicity measured by MTT assay. Each condition in quadruplicate across multiple experiments (see Table S2), error bars show standard deviation. **B.** Top - Lack of JK3/32 activity against replication (firefly luciferase activity) of HCV subgenomic replicon (JFH-1), bottom – efficient blockade of replicon replication using cyclosporine A (CsA). **C.** MRC PPU cellular kinase screen (see methods) testing 10 μM JK3/32, ~20x the IC_50_ concentration versus particle secretion.

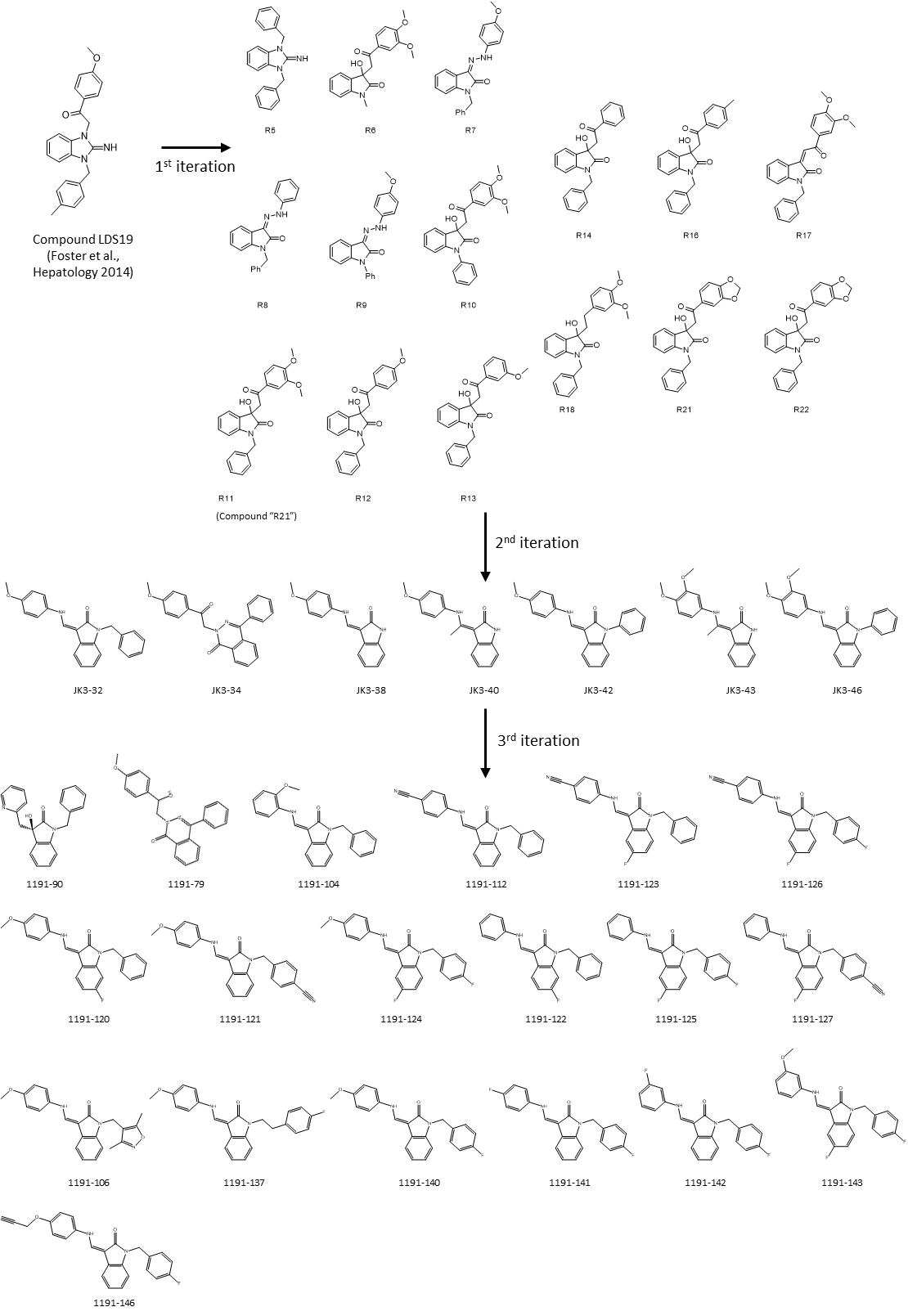

**Figure S3.** *Evolution of LDS19 compound series including JK3/32*. LDS19 was identified from an in silico screen using a 3ZD0-based channel structure model and demonstrated to have potent anti-p7 activity in vitro and in cell culture, as described previously(*24*). Three subsequent iterations of compound structure variations were undertaken to establish a comprehensive SAR and JK3/32 was established as the series lead. Compound RS11 (aka R21) was used as a negative control in cell entry assays due to a lack of activity versus genotype 1b and 3a HCV.

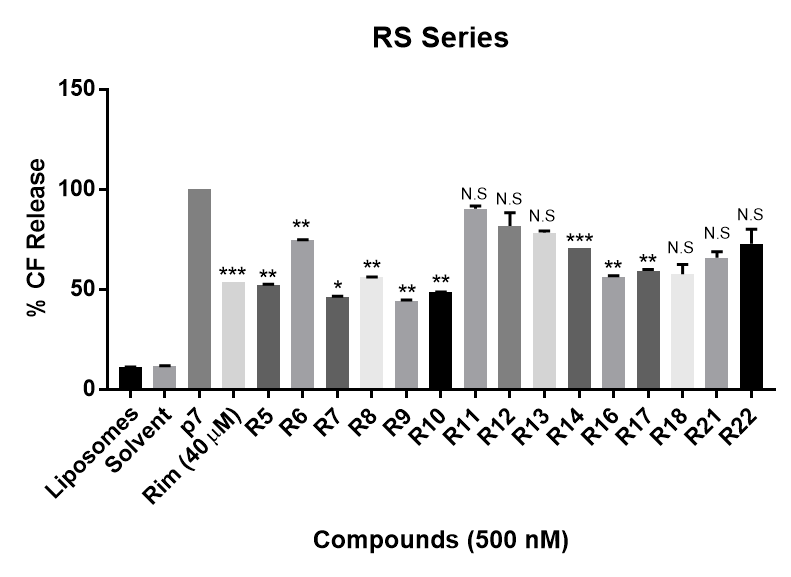

**Figure S4**. *In vitro dye release assay screen for activity of RS compounds versus genotype 1b p7*. RS series compounds were tested at a concentration of 500 nM versus recombinant J4 strain genotype 1b p7, alongside a positive control for inhibition (40 μM rimantadine). End-point fluorescence values from two experimental repeats were normalised against protein in the presence of no drug (100%). Error bars represent standard deviations (***p≤0.0001, **p≤0.001, *p≤0.05, Student T-Test, n=2).

**Figure S5*.*** *Effects of clinically advanced viroporin inhibitor, BIT225, versus particle secretion.* The amiloride derivative BIT225 was tested for effects against secreted HCV infectivity (J4/JFH-1 genotype 1b chimaera) using the optimised IncuCyte protocol. Conditions in quadruplicate, error bars show standard deviation. Calculated IC_50_ was ~17.7 μM, whereas CC50 was near identical at 18.6 μM.

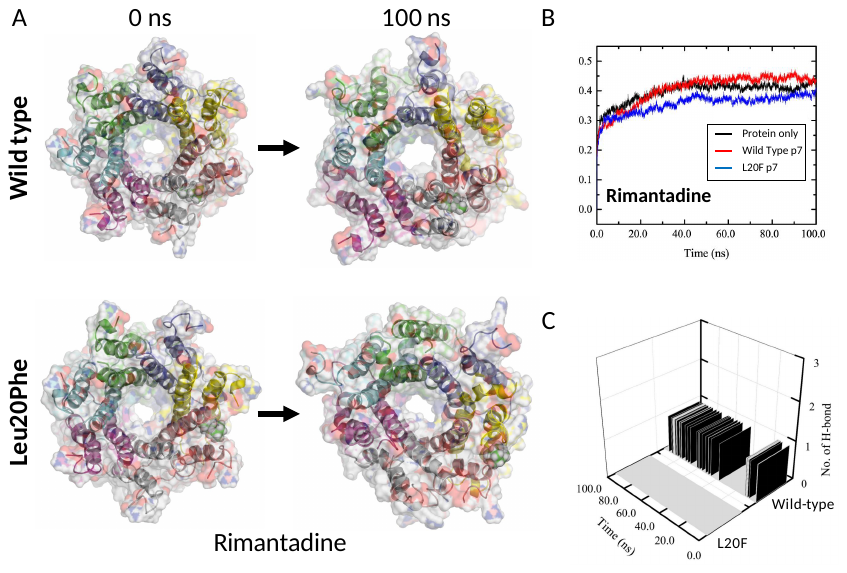

**Figure S6.** *Molecular Dynamic Simulation of rimantadine with heptameric 3ZD0-derived p7 and resistant Leu20Phe complexes*. An L20F polymorphism was introduced into the genotype 1b p7 sequence in the context of the 3ZD0 heptamer, then subjected to energy minimisation within a hydrated lipid bilayer. **A.** Location of rimantadine into the peripheral binding pocket led to stable, but weak interactions with wild type p7, whereas the Leu20Phe mutation caused a loss of binding interactions and migration of Rimantadine away from the pocket. This was also evident from RMSD traces (**B**), as well as the loss of H-bonds between Rimantadine and the protein (**C**).

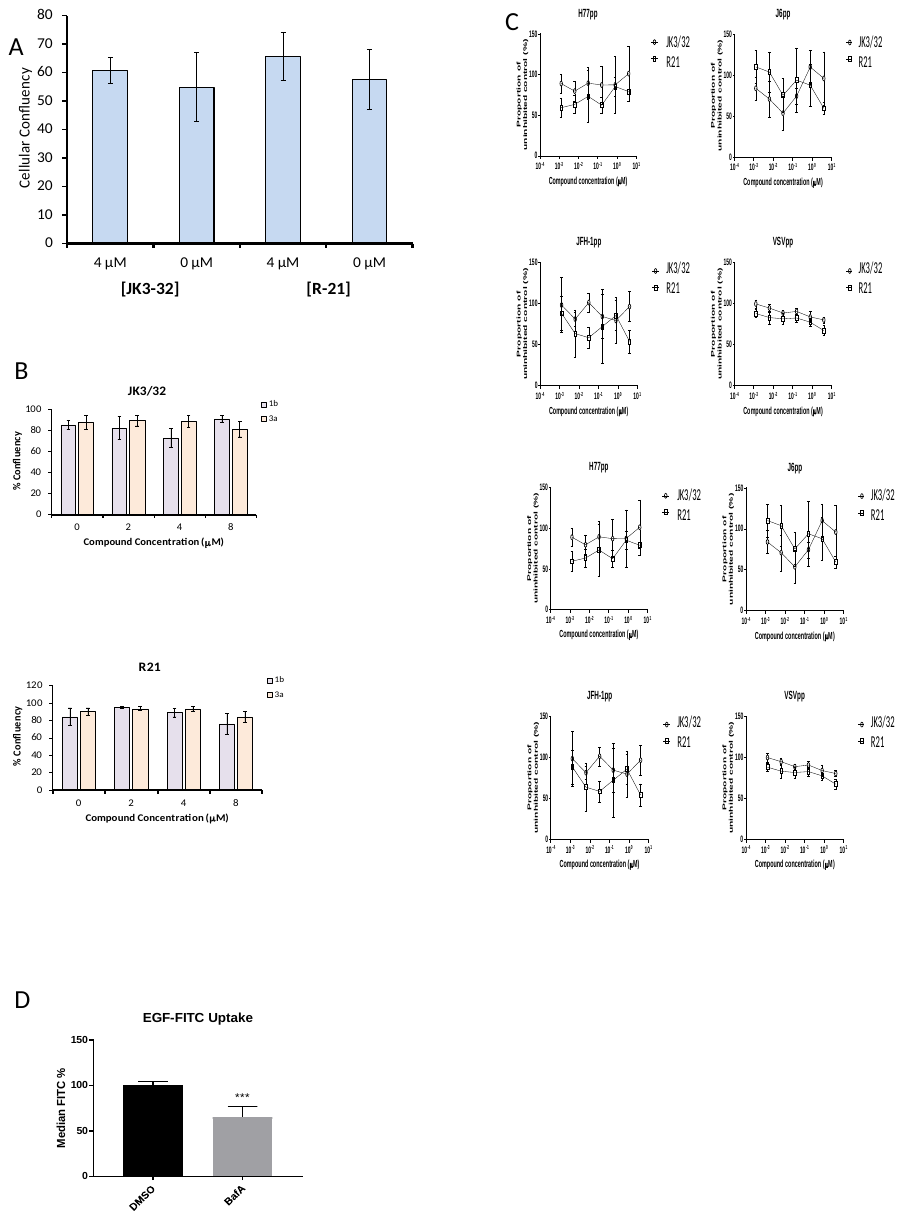

**Figure S7.** *Additional control experiments for JK3/32 effects during HCV entry*. **A, B**. Cell viability (Incucyte cell confluency) during treatment of Huh7 cells with JK3/32 or R21 during virus entry experiments shown in Fig 4a-c. **C.** Additional Lentiviral pseudotype assays using HCV envelopes derived from standard laboratory strains (H77, J6 and JFH-1), as well as a VSV-G control. **D.** Positive control for EGF uptake assays using 1 μM Bafilomycin A.

**Figure S8.** *IncuCyte image set from iodixinol gradients shown in figure 5*. 10 μL of each gradient fraction was added to 10 000 naïve Huh7 cells within each well of a black-walled 96-well plate. 48 h post-infection, cells were immunofluorescence stained for HCV NS5A protein and infectious units counted using the IncuCyte.

**Table S1.** *Consensus sequence and minority species for p7 amino acid positions based upon alignment of 1456 sequences from the EU HCV database*. JK3/32 binding site positions from the PDB:3ZD0-based model are highlighted in yellow.

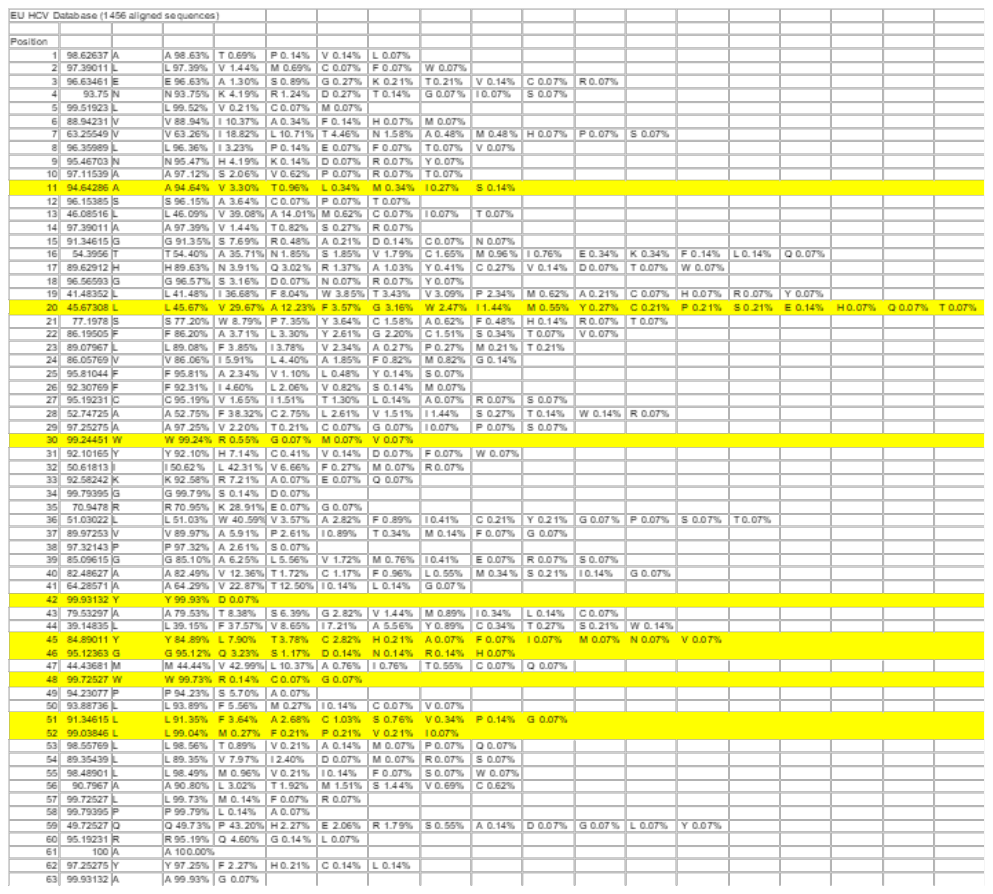

**Compound syntheses and purification**

***Synthesis of alexafluor-JK3/32 adduct (JK3/32-488):***

***3-[(dimethylamino)methylidene)]-2,3-dihydro-1H-indol-2-one*.**

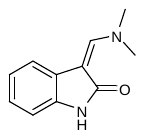

Dimethylformamide dimethylacetal (1.2 g, 10 mmol) was added with cooling to a suspension of oxindole (1.30 g, 9.80 mmol) in chloroform (15 mL) before the contents were stirred at room temperature for 15 mins, and heated under reflux for 4 h. The contents were cooled to room temperature and concentrated under reduced pressure to afford an orange solid, which was recrystallised from ethanol-diethyl ether to afford a yellow solid, which was used crude in the subsequent reaction.

***3-[(dimethylamino)methylidene]-1-{(4-fluorophenyl)methyl]-2,3-dihydro-1H-indol-2-one.***

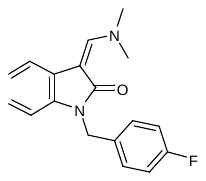

Caesium Carbonate (580 mg, 1.8 mmol) was added in a single portion with stirring to an acetonitrile (5 mL) suspension of 3-[(dimethylamino)methylidene)]-2,3-dihydro-1H-indol-2-one (200 mg, 1.1 mmol) at room temperature and the contents were stirred for a further 30 min, prior to the addition of 4-fluorobenzylbromide (300 mg, 1.6 mmol) *via* a syringe, after which the contents were stirred for 4 h at room temperature and then heated at 70 ^o^C for a further 1 h. The reaction mixture was cooled to room temperature, and transferred to a separating funnel, whereupon the reaction mixture was diluted with water and extracted into dichloromethane (2 x 15 ml). After drying over sodium sulphate, the contents were filtered and the organic phase concentrated under reduced pressure to yellow oil, which was chromatographed (SiO_2_; gradient elution; Hexane-EtOAc = 1 : 1 to 100 % EtOAc) to afford the desired product as pale yellow solid (210 mg, 67 %) which was carried forward to the next step without additional characterisation.

***(3Z)-1-[(4-fluorophenyl)methyl]-3-({[4-(prop-2-yn-1-yloxy)phenyl]amino}methylidene)-2,3-dihydro-1H-indol-2-one (1191-146)***

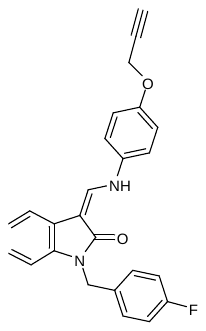

An ethanol (3 ml) suspension of 3-[(dimethylamino)methylidene]-1-{(4-fluorophenyl)methyl]-2,3-dihydro-1H-indol-2-one (53 mg, 0.18 mmol) was treated with 4-aminophenylpropargylether (35 mg, 0.23 mmol) and *p*-toluenesulphonic acid (52 mg, 0.28 mmol), and the contents were heated under reflux with stirring for 18 h until HPLC-MS had indicated consumption of the dimethylenamide. Upon gradual cooling to 60 ^o^C, together with intermittent scratching of the inside of the flask with a micro-spatula, there was obtained a precipitate that was filtered at 50 ^o^C, washed with cold ethanol and recrystallized from chloroform-diethyl ether to yield the desired product as a yellow solid (26 mg, 37%). IR: v_max_/cm^-1^ (solid): 3080, 1633, 1610. HPLC-MS: 2.24 min, 399.5 [M+H]^+^. ^1^H NMR (500 MHz, CDCl_3_): 10.64 (d, *J* = 13 Hz, 1H), 7.87 (d, *J* = 13 Hz, 1H), 7.30 (d, *J* = 7 Hz, 1H), 7.21 (br, 3H), 7.06 (d, *J* = 8.5 Hz, 2H), 6.93 (m, 4H), 6.71 (d, *J* = 8.5 Hz, 1H), 4.96 (s, 2H), 4.63 (d, *J* = 2.5 Hz, 2H), 2.47 (d, *J* = 2.5 Hz, 1H); ^13^C (300 MHz, CDCl_3_): 157.6, 149.5, 137.7, 132.8, 132.3, 129.2, 124.0, 121.3, 117.7, 116.4, 116.0, 115.7, 108.5, 99.6, 98.6, 78.4, 75.7, 56.3, 42.5.

***Alexafluor-JK3/32-triazole (JK3/32-488)*:**

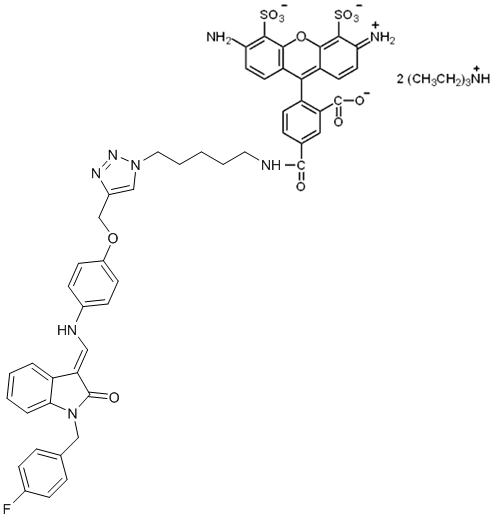

Alexafluor-488 (0.5 mg) was supplied in a light-proof centrifuge tube, to which was added a *tert*butanol (0.2 mL) suspension of the (3*Z*)-1-[(4-fluorophenyl)methyl]-3-({[4-(prop-2-yn-1-yloxy)phenyl]amino}methylidene)-2,3-dihydro-1H-indol-2-one (1191-146) (0.25 mg), followed by a solution (0.1 mL) aqueous ascorbic acid (0.35 mg). The contents were vortexed for 30 mins, before an aqueous solution (0.05 mL) of copper sulphate (0.20 mg) was added, and the contents shaken overnight at 40 ^o^C. Following this, the contents were concentrated on a Genevac, to a residue which was re-suspended in DMSO (0.20 mL) and centrifuged to disperse the solutes.

***Synthesis of JK3/32:***

***1-benzyl-3-[(dimethylamino)methylidene]-2,3-dihydro-1H-indol-2-one.***

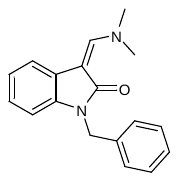

A DMF (4ml) suspension of (3*Z*)-3-[(dimethylamino)methylidene]-1,3-dihydro-2H-indol-2-one (300 mg, 1.6 mmol) was cooled to 0 ^o^C *via* an ice-bath with rapid stirring. Sodium hydride (125 mg, 60 % dispersion) was added in three portions over 10 min, and the resulting yellow suspension stirred for a further 20 min at 5 ^o^C, prior to the addition of benzyl bromide (330 mg, 1.90 mmol), and the contents left to stir for a further 45 min and allowed to warm to 25 ^o^C over this period. Saturated ammonium chloride solution was added dropwise with cooling, and the contents transferred with ethyl acetate to a separating funnel whereupon the organic phase was removed, washed with water and brine, and dried over sodium sulphate. Evaporation and chromatography (SiO_2_; gradient elution; hexane: EtOAc = 2 : 1 to 100 % EtOAc) afforded the desired product (210 mg, 47 %) as an oil which solidified upon standing. HPLC-MS: 1.89 min, 279.1 [M+H]^+^. IR: v_max_/cm^-1^ (solid): 1625: ^1^H NMR (500 MHz, CDCl_3_): 3.34–3.37, (6H, m, 2CH_3_), 5.06 (2H, s, CH_2_Ph), 6.74–7.42 (8H, m, ArH), 7.49 (1H, d, H-4), 7.68 (1H, s, CH).

***1-benzyl-3-[(4-methoxyphenyl)aminomethylidene]-2,3-dihydro-1H-indol-2-one (JK3/32)***

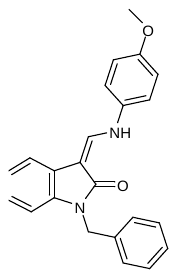

An ethanol (3 ml) solution of *p*-toluenesulfonic acid (26 mg, 0.14 mmol) was combined with 4-anisidine (17 mg, 0.14 mmol) at room temperature, and sonicated briefly to disperse the contents. 1-benzyl-3-[(dimethylamino)methylidene]-2,3-dihydro-1H-indol-2-one (35 mg, 0.13 mmol) was added in a single portion and the contents heated under reflux with stirring for 12 h to provide a precipitate, which was diluted with methanol (0.5 ml) and filtered at 50 ^o^C to afford the desired product as a yellow solid (18 mg, 39 %). IR: v_max_/cm^-1^ (solid): 3070, 1631, 1610. HPLC-MS: 2.21 min, 357.1[M+H]^+^. ^1^H NMR (500 MHz, CDCl_3_): δ 10.76 (d, *J* = 13 Hz, 1H), 7.97 (d, *J* = 13 Hz, 1H), 7.38 (d, *J* = 6.5 Hz, 1H), 7.31 (m, 3H), 7.24 (m, 2H), 7.12 (d, *J* = 8.5 Hz, 2H), 7.03 (m, 2H), 6.94 (d, *J* = 8.5 Hz, 2H), 6.82 (d, *J* = 8.0 Hz, 1H), 5.07 (s, 2H), 3.82 (s, 3H); ^13^C NMR (300 MHz, d_6_-DMSO) δ 167.8, 156.1, 139.4, 137.5, 137.1, 133.4, 128.6, 127.3, 127.0, 123.4, 120.7, 118.9, 117.8, 116.7, 115.1, 108.4, 97.4, 55.6, 42.3. HRMS (*m/z*): [M+H]^+^ calcd. for C_23_H_20_N_2_O_2_, 357.1598, found 357.1602.

***(3Z)-3-[(4-methoxyanilino)methylidene]-1-phenyl-1,3-dihydro-2H-indol-2-one (JK3-42)***

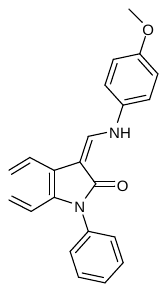

To *N*-phenyloxindole (0.9 mmol) was added dimethylformamide dimethyl acetal (5 mL) and the mixture heated at 70 ^o^C for 1 h. The mixture was concentrated, dissolved in ethanol (3 mL) and then *p*-toluenesulfonic acid (48 mg, 0.24 mmol) and 4-anisidine (30 mg, 0.24 mmol) added at room temperature and the contents heated under reflux with stirring for 12 h to provide a precipitate which was diluted with methanol (0.5 mL) and filtered at 50 ^o^C to afford the desired product as a pale yellow solid (19 mg, 42%). IR: v_max_/cm^-1^ (solid): 3080, 1633, 1610. HPLC-MS: 2.13 min, 345.1[M+H]^+^. ^1^H NMR (500 MHz, CDCl_3_): δ 10.78 (d, *J* = 13 Hz, 1H), 7.94 (d, *J* = 13 Hz, 1H), 7.33 (d, *J* = 6.9 Hz, 1H), 7.25 (m, 3H), 7.18 (m, 2H), 7.12 (d, *J* = 8.3 Hz, 2H), 7.03 (m, 2H), 6.94 (d, *J* = 8.3 Hz, 2H), 6.78 (d, *J* = 8.3 Hz, 1H), 3.87 (s, 3H); ^13^C NMR (300 MHz, d_6_-DMSO) δ 165.8, 158.1, 141.4, 138.5, 137.3, 135.1, 128.6, 127.6, 127.0, 125.9, 124.8, 123.1, 117.9, 116.7, 116.1, 109.3, 96.4, 55.7. HRMS (*m/z*): [M+H]^+^ calcd. for C_22_H_18_N_2_O_2_, 345.1575, found 345.1601.

***(3Z)-3-[(4-methoxyanilino)methylidene]-1,3-dihydro-2H-indol-2-one (JK3-38)***

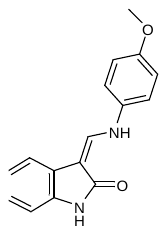

An ethanol (3 ml) solution of *p*-toluenesulfonic acid (26 mg, 0.14 mmol) was combined with 4-anisidine (17 mg, 0.14 mmol) at room temperature, and sonicated briefly to disperse the contents. (3*Z*)-3-[(dimethylamino)methylidene]-1,3-dihydro-2*H*-indol-2-one (0.14 mmol) was added in a single portion and the contents heated under reflux with stirring for 12 h to provide a precipitate, which was diluted with methanol (0.5 ml) and filtered at 50 ^o^C to afford the desired product as a yellow solid (15 mg, 39%). IR: v_max_/cm^-1^ (solid): 3100, 3050, 1640, 1610. HPLC-MS: 1.97 min, 266.1[M+H]^+^. ^1^H NMR (500 MHz, CDCl_3_): δ 10.50 (d, *J* = 11.0 Hz, 1H), 9.99 (s, 1H), 8.08 (d, *J* = 11.0 Hz, 1H), 7.55 (d, *J* = 6.9 Hz, 1H), 7.25 (m, 2H), 6.85-6.92 (4H, m), 6.78 (d, *J* = 8.3 Hz, 1H), 3.87 (s, 3H); ^13^C NMR (300 MHz, d_6_-DMSO) δ 165.8, 158.1, 141.4, 138.5, 135.1, 129.6, 126.0, 124.9, 124.0, 123.1, 117.5, 116.0, 115.1, 107.3, 93.4, 55.7. HRMS (*m/z*): [M+H]^+^ calcd. for C_16_H_14_N_2_O_2_, 266.1102, found 266.1197.

***(Z)-1-benzyl-3-(2-(4-methoxyphenyl)hydrazono)indolin-2-one (21-RS-7)***

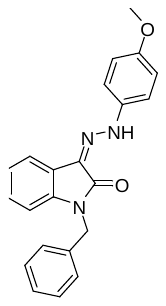

To a solution of *N*-benzyl isatin (0.05 g, 0.21 mmol) in ethanol (5 mL) was added *p*-methoxyphenylhydrazine hydrochloride (0.3 mL, 0.21 mmol) and the mixture heated at reflux for 2 h. The reaction mixture was cooled to ambient temperature and the resultant precipitate filtered and dried to give the title compound as a yellow powder (0.04 g, 0.11 mmol, 50%). IR: v_max_/cm^-1^ (solid): 1663, 1610. HPLC-MS (ES): 2.42 min, m/z = 737.6 (2M+Na)^+^ ^1^H NMR (300 MHz, DMSO-*d*_6_): δ 12.75 (s, 1H), 7.59 (dd, *J* = 7.2, 0.9 Hz 1H), 7.46 (d, *J* = 9.0 Hz, 2H), 7.38 – 7.25 (m, 5H), 7.22 (dd, *J* = 7.7, 1.3 Hz, 1H), 7.15 – 7.01 (m, 2H), 6.99 (d, *J* = 9.0 Hz, 2H), 5.03 (s, 2H), 3.76 (s, 3H) ppm; ^13^C NMR (300 MHz, CDCl_3_): δ 162.3, 156.2, 139.8, 136.4, 135.9, 128.8, 127.7, 127.4, 127.3, 125.6, 122.5, 121.6, 118.6, 115.7, 114.8, 109.2, 55.6, 43.2 ppm;; HRMS (*m/z*): [M+H]^+^ calcd for C_22_H_19_N_3_O_2_Na requires 380.1369 Found: 380.1374 (M+Na)^+^: v_max_/cm^-1^ (solid): 1663, 1610; M.pt: 145-147 °C.

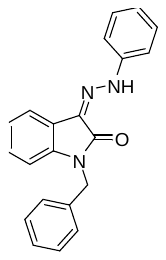
 ***(Z)-1-benzyl-3-(2-phenylhydrazono)indolin-2-one (21-RS-8):***

To a solution of *N*-benzyl isatin (0.05 g, 0.21 mmol) in ethanol (5 mL) was added phenylhydrazine hydrochloride (0.03 mg, 0.21 mmol) and the mixture heated at reflux for 2 h. The reaction mixture was cooled to ambient temperature and the resultant precipitate filtered and dried to give the title compound as a yellow powder (0.03 g, 0.09 mmol, 42%).

IR: v_max_/cm^-1^ (solid): 3060, 1663. ^1^H NMR (300 MHz, DMSO-*d*_6_): δ 12.72 (s, 1H), 7.64 – 7.59 (m, 1H), 7.52 – 7.46 (m, 2H), 7.43 – 7.30 (m, 6H), 7.30 – 7.22 (m, 2H), 7.16 – 7.07 (m, 1H), 7.10 – 7.01 (m, 2H), 5.03 (s, 2H) ppm; ^13^C NMR (300 MHz, DMSO-*d*_6_): 161.1, 142.4, 140.0, 136.3, 129.5, 128.7, 128.3, 127.5, 127.4, 126.5, 123.1, 122.5, 120.5, 118.5, 114.3, 109.8 ppm; HPLC-MS (ES): 2.07 min, m/z=677.7 (2M+Na)^+^; HRMS (*m/z*): [M+H]^+^ calcd for C_21_H_17_N_3_ONa requires 350.1264, Found: 350.1267 (M+Na)^+^,.; HPLC: RT = 4.51 min (100%); M.pt: 134-136 °C.

***(Z)-3-(2-(4-methoxyphenyl)hydrazono)-1-phenylindolin-2-one (21-RS-9):***

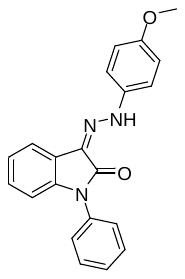

To a solution of *N*-phenyl isatin (0.05 g, 0.22 mmol) in ethanol (5 mL) was added *p-*methoxyphenylhydrazine hydrochloride (0.04 mg, 0.22 mmol) and the mixture heated at reflux for 2 h. The reaction mixture was cooled to ambient temperature and the resultant precipitate filtered and dried to give the title compound as a yellow powder (0.06 g, 0.17 mmol, 77%).

IR: v_max_/cm^-1^ (solid): 3186, 1673. ^1^H NMR (300 MHz, CDCl_3_): δ 12.88 (s, 1H), 7.69 – 7.60 (m, 1H), 7.52 – 7.45 (m, 2H), 7.43 – 7.33 (m, 4H), 7.26 (d, *J* = 9.0 Hz, 2H), 7.17 – 7.05 (m, 2H), 6.86 (d, *J* = 8.9 Hz, 2H), 3.75 (s, 3H) ppm; ^13^C NMR (300 MHz, CDCl_3_): δ 161.6, 156.2, 140.3, 136.3, 133.9, 129.6, 128.1, 127.4, 126.4, 125.3, 123.0, 121.6, 118.6, 115.7, 114.8, 109.7, 55.6 ppm; HPLC-MS (ES): : RT = 2.46 min, m/z= 709.9 (2M+Na)^+^; HRMS (*m/z*): [M+H]^+^ calcd for C_21_H_17_N_3_O_2_Na requires 366.1209: 366.1209 (M+Na)^+^,;;; M.pt: 195-197 °C.

***1-benzyl-3-(2-(3,4-dimethoxyphenyl)-2-oxoethyl)-3-hydroxyindolin-2-one (21-RS-11 (aka “R21”)):***

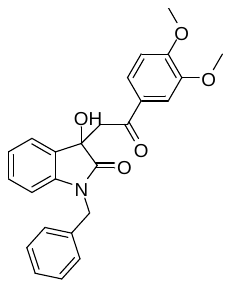

To a microwave vial was charged *N*-benzyl isatin (0.20 g, 0.84 mmol), 3,4-dimethoxyacetophenone (0.17 g, 0.93 mmol), diethylamine (2 drops) and ethanol (2 mL). The reaction was heated at 100 °C and 300 W within a microwave reactor for 1-2 h. The crude product was purified *via* column chromatography (50:50 hexane:ethyl acetate) to yield the title compound as a cream powder (0.24 g, 0.58 mmol, 68%). IR: v_max_/cm^-1^ (solid): 3329, 1668, 1598. ^1^H NMR (300 MHz, CDCl_3_): δ 7.54 (d, *J* = 8.4 Hz, 1H), 7.48 (s, 1H), 7.43 (d, *J* = 7.4 Hz), 7.40 – 7.29 (m, 5H), 7.21 (t, *J* = 7.8 Hz, 1H), 7.01 (t, *J* = 7.5 Hz, 1H), 6.86 (d, *J* = 8.4 Hz, 1H), 6.74 (d, *J* = 7.8 Hz, 1H), 4.95 (s, 2H), 4.83 (br s, 1H), 3.95 (s, 3H), 3.91 (s, 2H), 3.86 (d, *J* = 17.6 Hz, 1H), 3.59 (d, *J* = 17.1 Hz, 1H) ppm; ^13^C NMR (300 MHz, CDCl_3_): δ 196.8, 176.6, 153.9, 149.0, 142.8, 135.5, 130.2, 129.8, 129.6, 128.8, 127.7, 127.3, 124.1, 123.3, 123.1, 110.0, 109.9, 109.7, 80.5, 74.7, 56.2, 56.0, 43.9 ppm; HPLC-MS (ES): RT = 1.87 min, m/z = 857.6 (2M+Na)^+^; HRMS (*m/z*): [M+H]^+^ calcd for C_25_H_23_NO_5_Na requires 440.1468, Found: 440.1488 (M+Na)^+^. M.pt: 157-159 °C.

***1-benzyl-3-(2-(3,4-dimethoxyphenyl)-2-oxoethylidene)indolin-2-one (21-RS-17)***

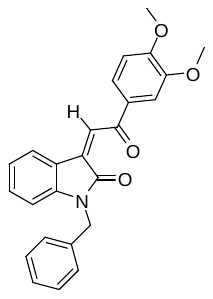

To a stirred solution of 1-benzyl-3-(2-(3,4-dimethoxyphenyl)-2-oxoethyl)-3-hydroxyindolin-2-one (21-RS-11) (0.04 g, 0.08 mmol) in acetic acid (5 mL) was added 12M aqueous hydrochloric acid (2 mL). The mixture was heated at 80 °C for 0.5 h then stirred at ambient temperature for 2 days. The mixture was quenched with cold aqueous sodium bicarbonate (20 mL), extracted with ethyl acetate (3x20 mL) and concentrated to dryness. The crude product was purified *via* column chromatography (50:50 hexane:ethyl acetate) to give the title compound as an orange powder (0.02 g, 0.05 mmol, 57%). IR: v_max_/cm^-1^ (solid): 1705, 1599. ^1^H NMR (300 MHz, CDCl_3_): δ 8.19 (d, *J* = 7.6 Hz, 1H), 7.88 (s, 1H), 7.72 (dd, *J* = 8.4, 2.0 Hz, 1H), 7.61 (d, *J* = 2.0 Hz, 1H), 7.31 – 7.12 (m, 6H), 6.96 – 6.83 (m, 2H), 6.64 (d, *J* = 7.8 Hz, 1H), 4.91 (s, 2H), 3.92 (d, *J* = 2.4 Hz, 6H) ppm; ^13^C NMR (300 MHz, CDCl_3_): δ 189.6, 168.2, 154.2, 149.5, 144.9, 135.6, 135.5, 132.2, 130.8, 128.9, 127.8, 127.6, 127.3, 127.2, 124.4, 122.8, 110.1, 109.2, 105.0, 80.2, 56.2, 56.1, 43.9 ppm; HPLC-MS (ES): RT = 2.18 min, m/z = 00.1 (M+H)^+^; HRMS (*m/z*): [M+H]^+^ calcd for C_25_H_21_NO_4_Na requires 422.1363 Found: 422.1356 (M+Na)^+^, ; M.pt: 146-148 °C.

***(3Z)-1-benzyl-3-[(2-methoxyanilino)methylidene]-1,3-dihydro-2H-indol-2-one (1191-104)***

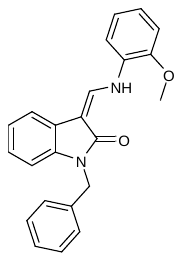

An ethanol (3 mL) solution of *p*-toluenesulfonic acid (52 mg, 0.27 mmol) was combined with 4-anisidine (34 mg, 0.27 mmol) at room temperature, and sonicated briefly to disperse the contents. 1-benzyl-3-[(dimethylamino)methylidene]-2,3-dihydro-1H-indol-2-one (68 mg, 0.27 mmol) was added in a single portion and the contents heated under reflux with stirring for 12 h to provide a precipitate, which was diluted with methanol (0.5 mL) and filtered at 50 ^o^C to afford the desired product as a yellow solid (70 mg, 75%). IR: v_max_/cm^-1^ (solid): 3079, 1630. HPLC-MS: 2.28 min, 357.1 [M+H]^+^. ^1^H NMR (500 MHz, CDCl_3_): δ 11.02 (d, *J* = 12.5 Hz, 1H), 8.10 (d, *J* = 12.5 Hz, 1H), 7.43 (d, *J* = 8.5 Hz, 1H), 7.32 (m, 6H), 7.05 (m, 4H), 6.99 (d, *J* = 8.0 Hz, 1H), 6.80 (d, *J* = 7.8 Hz, 1 H), 5.12 (s, 2H), 4.09 (s, 3H); ^13^C NMR (125 MHz, CDCl_3_) 157.6, 149.1, 132.7, 131.7, 130.4, 125.8, 123.5, 122.0, 119.2, 118.3, 115.7, 111.8, 110.8, 107.5, 105.9, 103.8, 97.2, 50.7, 37.8.

***4-{[(Z)-(1-benzyl-2-oxo-1,2-dihydro-3H-indol-3-ylidene)methyl]amino}benzonitrile (1191-112)***

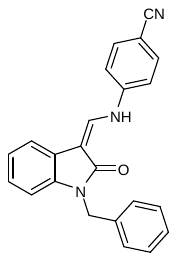

An ethanol (3 mL) solution of *p*-toluenesulfonic acid (52 mg, 0.27 mmol) was combined with 4-aminophenylnitrile (33 mg, 0.27 mmol) at room temperature. 1-benzyl-3-[(dimethylamino)methylidene]-2,3-dihydro-1H-indol-2-one (68 mg, 0.27 mmol) was added in a single portion and the contents heated under reflux with stirring for 12 h to provide a precipitate, which was diluted with methanol (0.5 mL) and filtered at 50 ^o^C to afford the desired product as a yellow solid (50 mg, 52%). IR: v_max_/cm^-1^ (solid): 3079, 2127, 1630. HPLC-MS: 2.20 min, 352.5 [M+H]^+^. ^1^H NMR (500 MHz, CDCl_3_): δ 10.89 (d, *J* = 13.5 Hz, 1H), 7.89 (d, J = 12.8 Hz, 1H), 7.58 (d, *J* = 7.5 Hz, 2H), 7.35 (d, *J* = 9.0 Hz, 1H), 7.24 (m, 4H), 7.18 (m, 1H), 7.12 (d, *J* = 7.5 Hz, 2H), 7.04 (t, *J* = 7.5 Hz, 1H), 6.97 (t, *J* = 6.0 Hz, 1H), 6.75 (d, *J* = 6.0 Hz, 1H), 4.92 (s, 2H); ^13^C NMR (125 MHz, CDCl_3_) 153.8, 143.6, 134.1, 130.8, 128.9, 127.7, 127.2, 125.7, 122.6, 121.8, 118.6, 116.9, 115.1, 109.16, 105.7, 104.6, 93.1, 43.4.

***(3Z)-1-benzyl-6-fluoro-3-[(4-methoxyanilino)methylidene]-1,3-dihydro-2H-indol-2-one (1191-121)***

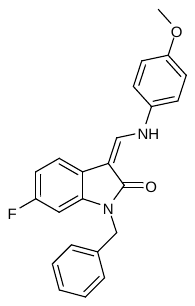

A DMF (4ml) suspension of (3*Z*)-3-[(dimethylamino)methylidene]-6-fluoro-1,3-dihydro-2*H*-indol-2-one (300 mg, 1.60 mmol) was cooled to 0 ^o^C *via* an ice-bath with rapid stirring. Sodium hydride (125 mg, 60% dispersion) was added in three portions over 10 min, and the resulting yellow suspension stirred for a further 20 min at 5 ^o^C, prior to the addition of benzyl bromide (1.80 mmol) and the contents left to stir for a further 45 min and allowed to warm to 25 ^o^C over this period. Saturated ammonium chloride solution was added dropwise with cooling, and the contents transferred with ethyl acetate to a separating funnel whereupon the organic phase was removed, washed with water and brine, and dried over sodium sulphate. Evaporation and chromatography (SiO_2_; gradient elution; hexane : EtOAc = 2 : 1 to 100 % EtOAc) afforded the desired product as an oil which solidified upon standing. This was then added to an ethanol (3 mL) solution of *p*-toluenesulfonic acid (52 mg, 0.27 mmol) and 4-anisidine (34 mg, 0.27 mmol) and the contents heated under reflux with stirring for 12 h to provide a precipitate, which was diluted with methanol (0.5 mL) and filtered at 50 ^o^C to afford the desired product as a pale yellow solid (45 mg, 48%). IR: v_max_/cm^-1^ (solid): 3099, 1630. HPLC-MS: 2.24 min, 375.3 [M+H]^+^. ^1^H NMR (500 MHz, CDCl_3_): δ 10.79 (d, *J* = 13.5 Hz, 1H), 7.92 (d, *J* = 13.5 Hz, 1H), 7.28 (m, 5H), 7.12 (d, *J* = 10 Hz, 2H), 7.07 (m, 1H), 6.93 (d, *J* = 8.5 Hz, 2H), 6.70 (m, 2H), 5.05 (s, 2H), 3.82 (s, 3H); ^13^C NMR (300 MHz, CDCl_3_): 155.4, 148.1, 138.6, 133.5, 132.2, 128.7, 127.7, 127.4, 127.2, 126.0, 125.2, 118.1, 115.3, 109.9, 108.9, 103.3, 99.2, 55.8, 43.8.

***4-({(3Z)-5-fluoro-3-[(4-methoxyanilino)methylidene]-2-oxo-2,3-dihydro-1H-indol-1-yl}methyl)benzonitrile (1191-120)***

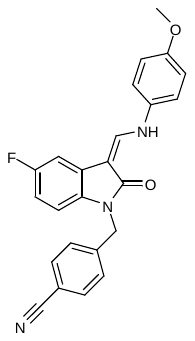

A DMF (4ml) suspension of (3*Z*)-3-[(dimethylamino)methylidene]-5-fluoro-1,3-dihydro-2*H*-indol-2-one (300 mg, 1.6 mmol) was cooled to 0 ^o^C *via* an ice-bath with rapid stirring. Sodium hydride (125 mg, 60% dispersion) was added in three portions over 10 min, and the resulting yellow suspension stirred for a further 20 min at 5^o^C, prior to the addition of 4-cyanophenylmethyl bromide (369 mg, 1.90 mmol) and the contents left to stir for a further 45 min and allowed to warm to 25 ^o^C over this period. Saturated ammonium chloride solution was added dropwise with cooling, and the contents transferred with ethyl acetate to a separating funnel whereupon the organic phase was removed, washed with water and brine, and dried over sodium sulfate. Evaporation and chromatography (SiO_2_; gradient elution; hexane : EtOAc = 2 : 1 to 100 % EtOAc) afforded the desired product as an oil. This was then added to an ethanol (3 mL) solution of *p*-toluenesulfonic acid (52 mg, 0.27 mmol) and 4-anisidine (34 mg, 0.27 mmol) at room temperature and the contents heated under reflux with stirring for 12 h to provide a precipitate which was diluted with methanol (0.5 mL) and filtered at 50 ^o^C to afford the desired product as a yellow solid (51 mg, 48%). IR: v_max_/cm^-1^ (solid): 3130, 2145, 1620. HPLC-MS: 2.02 min, 400.4 [M+H]^+^. ^1^H NMR (500 MHz, CDCl_3_): δ 10.75 (d, *J* = 13 Hz, 1H), 7.95 (d, *J* = 13 Hz, 1H), 7.61 (d, *J* = 10 Hz, 2H), 7.38 (d, *J* = 8.5 Hz, 2H), 7.12 (d, *J* = 10 Hz, 2H), 7.09 (m, 1H), 6.94 (d, *J* = 8.5 Hz, 2H), 6.73 (t, *J* = 7.5 Hz, 1H), 6.60 (dd, *J* = 8.5 Hz, 4.2 Hz, 1H), 5.1 (s, 2H), 3.82 (s, 3H); ^13^C NMR (300 MHz, CDCl_3_) 169.1, 160.3, 157.0, 142.2, 139.1, 133.1, 132.9, 127.9, 118.7, 118.1, 115.2, 111.6, 110.4, 108.6, 55.9, 43.1.

***(3Z)-5-fluoro-1-[(4-fluorophenyl)methyl]-3-[(4-methoxyanilino)methylidene]-1,3-dihydro-2H-indol-2-one (1191-124)***

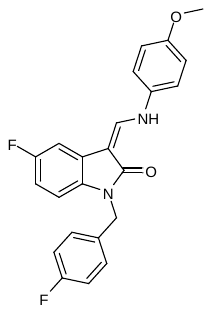

A DMF (4ml) suspension of (3*Z*)-3-[(dimethylamino)methylidene]-5-fluoro-1,3-dihydro-2*H*-indol-2-one (300 mg, 1.6 mmol) was cooled to 0 ^o^C. Sodium hydride (125 mg, 60% dispersion) was added in three portions over 10 min, and the resulting yellow suspension stirred for a further 20 min at 5 ^o^C, prior to the addition of 4-fluorophenylmethyl bromide (1.80 mmol), and the contents left to stir for a further 45 min. Saturated ammonium chloride solution was added dropwise with cooling, and the contents transferred with ethyl acetate to a separating funnel whereupon the organic phase was removed, washed with water and brine, and dried over sodium sulphate. Evaporation and chromatography (SiO_2_; gradient elution; hexane : EtOAc = 2 : 1 to 100 % EtOAc) afforded the desired intermediate as an oil. This was then added to an ethanol (3 mL) solution of *p*-toluenesulfonic acid (52 mg, 0.27 mmol) and 4-anisidine (34 mg, 0.27 mmol) at room temperature and the contents heated under reflux with stirring for 12 h to provide a precipitate which was diluted with methanol (0.5 mL) and filtered at 50 ^o^C to afford the desired product as a yellow solid (51 mg, 53%). IR: v_max_/cm^-1^ (solid): 3115, 1625. HPLC-MS: 2.26 min, 393.2 [M+H]^+^. ^1^H NMR (500 MHz, CDCl_3_): δ 10.71 (d, *J* = 13.5 Hz, 1H), 7.86 (d, *J* = 13.0 Hz, 1H), 7.19 (br, 1H), 7.05 (d, *J* = 8.5 Hz, 2H), 6.99 (d, *J* = 8.5 Hz, 1H), 6.92 (m, 3H), 6.87 (d, *J* = 7.7 Hz, 2H), 6.66 (t, *J* = 7.7 Hz, 1H), 6.59 (br, 1H), 4.93 (s, 2H), 3.76 (s, 3H); ^13^C NMR (300 MHz, CDCl_3_): 157.9, 149.6, 140.2, 138.0, 133.4, 133.2, 132.9, 128.9, 122.2, 118.0, 115.8, 115.6, 115.2, 110.2, 108.8, 103.4, 98.1, 55.6, 42.6.

***(3Z)-1-[(3,5-dimethyl-1,2-oxazol-4-yl)methyl]-3-[(4-methoxyanilino)methylidene]-1,3-dihydro-2H-indol-2-one (1191-106)***

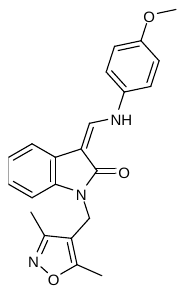

A DMF (4mL) suspension of (3*Z*)-3-[(dimethylamino)methylidene]-1,3-dihydro-2*H*-indol-2-one (300 mg, 1.60 mmol) was cooled to 0 ^o^C *via* an ice-bath with rapid stirring. Sodium hydride (125 mg, 60% dispersion) was added in three portions over 10 min, and the resulting yellow suspension stirred for a further 20 min at 5 ^o^C, prior to the addition of 2,5-dimethylisoxazole-4-methylchloride (1.80 mmol) and the contents left to stir for a further 45 min. Saturated ammonium chloride solution was added dropwise with cooling, and the contents transferred with ethyl acetate to a separating funnel whereupon the organic phase was removed, washed with water and brine, and dried over sodium sulphate. Evaporation and chromatography (SiO_2_; gradient elution; hexane : EtOAc = 2 : 1 to 100 % EtOAc) afforded an intermediate as an oil. This was then added to an ethanol (3 mL) solution of *p*-toluenesulfonic acid (48 mg, 0.24 mmol) and 4-anisidine (30 mg, 0.24 mmol) at room temperature and the contents heated under reflux with stirring for 12 h to provide a precipitate which was diluted with methanol (0.5 mL) and filtered at 50 ^o^C to afford the desired product as a pale yellow solid (57 mg, 54%). IR: v_max_/cm^-1^ (solid): 3120, 1625. HPLC-MS: 1.91 min, 375.3 [M+H]^+^. ^1^H NMR (500 MHz, CDCl_3_): δ 10.67 (d, *J* = 13 Hz, 1H), 7.97 (d, *J* = 13 Hz, 1H), 7.72 (d, *J* = 10 Hz, 1H), 7.39 (d, *J* = 10 Hz, 1H), 7.15 (d, *J* = 10 Hz, 1H), 7.08 (m, 2H), 6.96 (d, *J* = 10 Hz, 2H), 6.74 (d, *J* = 8 Hz, 1H), 4.84 (s, 2H), 3.86 (s, 3H), 2.42 (s, 3H), 2.25 (s, 3H); ^13^C NMR (300 MHz, CDCl_3_) 168.9, 166.9, 159.5, 156.7, 138.4, 137.0, 133.6, 129.0, 126.3, 123.9, 121.7, 117.9, 116.1, 115.3, 109.7, 108.5, 55.8, 32.5, 11.4, 10.6.

***(3Z)-3-[(4-methoxyanilino)methylidene]-1-(2-phenylethyl)-1,3-dihydro-2H-indol-2-one (1191-137)***

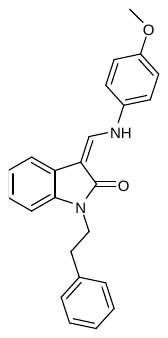

A DMF (4mL**)** suspension of (3*Z*)-3-[(dimethylamino)methylidene]-1,3-dihydro-2*H*-indol-2-one (300 mg, 1.6 mmol) was cooled to 0 ^o^C *via* an ice-bath with rapid stirring. Sodium hydride (125 mg, 60% dispersion) was added in three portions over 10 min, and the resulting yellow suspension stirred for a further 20 min at 5 ^o^C, prior to the addition of phenylethyl bromide (369 mg, 1.90 mmol), and the contents left to stir for a further 45 min and allowed to warm to 25 ^o^C over this period. Saturated ammonium chloride solution was added dropwise with cooling, and the contents transferred with ethyl acetate to a separating funnel whereupon the organic phase was removed, washed with water and brine, and dried over sodium sulphate. Evaporation and chromatography (SiO_2_; gradient elution; hexane : EtOAc = 2 : 1 to 100 % EtOAc) afforded an intermediate product as an oil which solidified upon standing. The material was used immediately in the subsequent reaction. The oil was added to an ethanol (3 mL) solution of *p*-toluenesulfonic acid (53 mg, 0.27 mmol) and 4-anisidine (35 mg, 0.27 mmol) at room temperature and the contents heated under reflux with stirring for 12 h to provide a precipitate which was diluted with methanol (0.5 mL) and filtered at 50 ^o^C to afford the desired product as a pale yellow solid (49 mg, 44%). IR: v_max_/cm^-1^ (solid): 3095, 1620.HPLC-MS: 2.29 min, 371.2 [M+H]^+^. ^1^H NMR (500 MHz, CDCl_3_): δ 10.61 (d, *J* = 13 Hz, 1H), 7.86 (d, *J* = 13 Hz, 1H), 7.30 (d, *J* = 8.0 Hz, 1H), 7.22 (m, 3H), 7.17 (m, 2H), 7.03 (m, 3H), 6.99 (d, *J* = 7.0 Hz, 1H), 6.85 (d, *J* = 7.0 Hz, 2H), 6.81 (d, *J* = 8.0 Hz, 1H), 3.99 (t, *J* = 7.5 Hz, 2H), 3.77 (s, 3H), 2.95 (t, *J* = 7.5 Hz, 2H); ^13^C NMR (300 MHz, CDCl_3_) 166.2, 149.5, 137.6, 133.7, 133.1, 131.4, 129.6, 128.9, 128.6, 128.4, 127.7, 121.1, 117.6, 117.5, 115.1, 108.4, 98.5, 55.8, 34.7, 33.5.

***(3Z)-1-[(4-fluorophenyl)methyl]-3-[(4-methoxyanilino)methylidene]-1,3-dihydro-2H-indol-2-one (1191-140)***

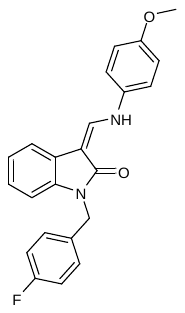

A DMF (4mL) suspension of (3*Z*)-3-[(dimethylamino)methylidene]-1,3-dihydro-2*H*-indol-2-one (300 mg, 1.6 mmol) was cooled to 0 ^o^C *via* an ice-bath with rapid stirring. Sodium hydride (125 mg, 60% dispersion) was added in three portions over 10 min, and the resulting yellow suspension stirred for a further 20 min at 5 ^o^C, prior to the addition of 4-flurophenylbromide (1.65 mmol), and the contents left to stir for a further 45 min. Saturated ammonium chloride solution was added dropwise and the contents transferred with ethyl acetate to a separating funnel whereupon the organic phase was removed, washed with water and brine, and dried over sodium sulphate. Evaporation and chromatography (SiO_2_; gradient elution; hexane : EtOAc = 2 : 1 to 100 % EtOAc) afforded an oil which was added to ethanol (3 mL) and *p*-toluenesulfonic acid (53 mg, 0.27 mmol) and 4-anisidine (35 mg, 0.27 mmol) at room temperature and the contents heated under reflux with stirring for 12 h to provide a precipitate which was diluted with methanol (0.5 mL) and filtered at 50 ^o^C to afford the desired product as a pale yellow solid (67 mg, 63%). IR: v_max_/cm^-1^ (solid): 3105, 1635.HPLC-MS: 2.24 min, 375.3 [M+H]^+^. ^1^H NMR (500 MHz, CDCl_3_): δ 10.63 (d, *J* = 13 Hz, 1H), 7.87 (d, *J* = 13 Hz, 1H), 7.31 (d, *J* = 7.5 Hz, 1H), 7.21 (m, 2H), 7.04 (d, *J* = 9 Hz, 2H), 6.94 (m, 3H), 6.85 (d, *J* = 8.5 Hz, 2H), 6.72 (d, *J* = 8.0 Hz, 1H), 4.97 (s, 2H), 3.75 (s, 3H); ^13^C NMR (300 MHz, CDCl_3_) 165.1, 156.5, 137.9, 137.4, 136.8, 135.3, 133.5, 132.6, 130.3, 128.8, 123.9, 123.8, 121.4, 117.1, 115.6, 115.1, 108.7, 98.1, 55.7, 42.7.

***(3Z)-3-[(4-fluoroanilino)methylidene]-1-[(4-fluorophenyl)methyl]-1,3-dihydro-2H-indol-2-one (****1191-141)*

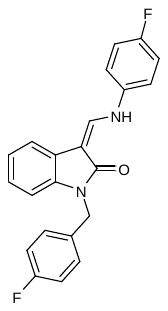

A DMF (4ml) suspension of (3*Z*)-3-[(dimethylamino)methylidene]-1,3-dihydro-2*H*-indol-2-one (300 mg, 1.6 mmol) was cooled to 0 ^o^C *via* an ice-bath with rapid stirring. Sodium hydride (125 mg, 60 % dispersion) was added in three portions over 10 min, and the resulting yellow suspension stirred for a further 20 min at 5 ^o^C, prior to the addition of 4-flurophenylbromide (1.65 mmol), and the contents left to stir for a further 45 min. Saturated ammonium chloride solution was added dropwise and the contents transferred with ethyl acetate to a separating funnel whereupon the organic phase was removed, washed with water and brine, and dried over sodium sulphate. Evaporation and chromatography (SiO_2_; gradient elution; hexane : EtOAc = 2 : 1 to 100 % EtOAc) afforded the desired product as an oil which was used immediately in the next reaction. The oil was added to ethanol (3 mL) and *p*-toluenesulfonic acid (53 mg, 0.27 mmol) and 4-fluorophenylaniline (0.27 mmol) at room temperature and the contents heated under reflux with stirring for 12 h to provide a precipitate which was diluted with methanol (0.5 mL) and filtered at 50 ^o^C to afford the desired product as a pale yellow solid (45 mg, 57%). IR: v_max_/cm^-1^ (solid): 3130, 1615. HPLC-MS: 2.20 min, 362.4 [M+H]^+^. ^1^H NMR (500 MHz, CDCl_3_): δ 10.60 (d, *J* = 12 Hz, 1H), 7.99 (d, *J* = 12 Hz, 1H), 7.23-7.39 (m, 3H), 7.15 (d, *J* = 9 Hz, 2H), 6.91 (m, 3H), 6.80 (d, *J* = 8.5 Hz, 2H), 6.77 (d, *J* = 7.9 Hz, 1H), 4.90 (s, 2H); ^13^C NMR (300 MHz, CDCl_3_) 165.1, 156.5, 139.1, 138.1, 136.7, 136.1, 133.5, 132.4, 131.1, 128.9, 124.0, 122.8, 121.8, 117.0, 114.1, 111.1, 104.6, 97.5, 58.9

***(3Z)-5-fluoro-3-[(anilino)methylidene]-1-[(4-fluorophenyl)methyl]-1,3-dihydro-2H-indol-2-one (****1191-125)*

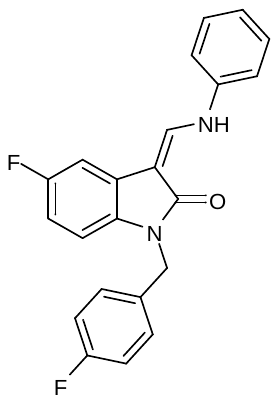

A DMF (4ml) suspension of (3*Z*)-3-[(dimethylamino)methylidene]-5-fluoro-1,3-dihydro-2*H*-indol-2-one (300 mg, 1.6 mmol) was cooled to 0 ^o^C *via* an ice-bath with rapid stirring. Sodium hydride (125 mg, 60% dispersion) was added in three portions over 10 min, and the resulting yellow suspension stirred for a further 20 min at 5 ^o^C, prior to the addition of 4-fluorobenzyl lbromide (1.65 mmol), and the contents left to stir for a further 45 min. Saturated ammonium chloride solution was added dropwise and the contents transferred with ethyl acetate to a separating funnel whereupon the organic phase was removed, washed with water and brine, and dried over sodium sulphate. Evaporation and chromatography (SiO_2_; gradient elution; hexane : EtOAc = 2 : 1 to 100 % EtOAc) afforded an oil which was used immediately in the next reaction. The oil was added to ethanol (3 mL) and *p*-toluenesulfonic acid (53 mg, 0.27 mmol) and aniline (0.27 mmol) at room temperature and the contents heated under reflux with stirring for 12 h to provide a precipitate which was diluted with methanol (0.5 mL) and filtered at 50 ^o^C to afford the desired product as a pale yellow solid (43 mg). IR: v_max_/cm^-1^ (solid): 3130, 1620. HPLC-MS: 2.30 min, 362.1 [M+H]^+^. ^1^H NMR (500 MHz, CDCl_3_): δ 10.65 (d, *J* = 11.5 Hz, 1H), 7.99 (d, *J* = 11.5 Hz, 1H), 7.15-7.39 (m, 6H), 6.91-7.12 (m, 3H), 6.77 (m, 3H), 4.90 (s, 2H).

***4-{[(Z)-(1-(4-fluoro)phenylmethyl-2-oxo-1,2-dihydro-5-fluoro-3H-indol-3-ylidene)methyl]amino}benzonitrile (****1191-126)*

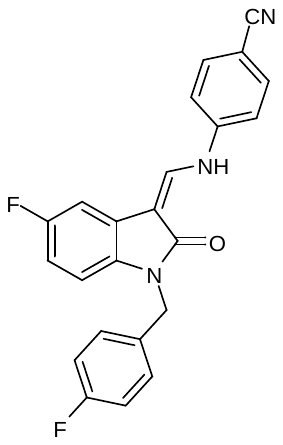

A DMF (4ml) suspension of (3*Z*)-3-[(dimethylamino)methylidene]-5-fluoro-1,3-dihydro-2*H*-indol-2-one (300 mg, 1.6 mmol) was cooled to 0 ^o^C *via* an ice-bath with rapid stirring. Sodium hydride (125 mg, 60% dispersion) was added in three portions over 10 min, and the resulting yellow suspension stirred for a further 20 min at 5 ^o^C, prior to the addition of 4-fluorobenzylbromide (1.65 mmol), and the contents left to stir for a further 45 min. Saturated ammonium chloride solution was added dropwise and the contents transferred with ethyl acetate to a separating funnel whereupon the organic phase was removed, washed with water and brine, and dried over sodium sulfate. Evaporation and chromatography (SiO_2_; gradient elution; hexane : EtOAc = 2 : 1 to 100 % EtOAc) afforded an oil which was used immediately in the next reaction. The oil was added to ethanol (3 mL) and *p*-toluenesulfonic acid (53 mg, 0.27 mmol) and aniline (0.27 mmol) at room temperature and the contents heated under reflux with stirring overnight to provide a precipitate which was diluted with methanol (2 x 0.5 mL) and filtered at 50 ^o^C to afford the desired product as a yellow solid (57 mg). IR: v_max_/cm^-1^ (solid): 3100, 2200, 1615. HPLC-MS: 2.00 min, 387.1 [M+H]^+^. ^1^H NMR (500 MHz, CDCl_3_): δ 10.71 (d, *J* = 12.0 Hz, 1H), 7.86 (d, *J* = 12.0 Hz, 1H), 7.45 (br, 1H), 7.35 (d, *J* = 8.5 Hz, 2H), 7.15 (d, *J* = 8.5 Hz, 1H), 6.92-6.99 (m, 3H), 6.83 (m, 3H), 6.76 (t, *J* = 7.5 Hz, 1H), 4.90 (s, 2H).
